# Discovery of Potent Inhibitors of α-Synuclein Aggregation Using Structure-Based Iterative Learning

**DOI:** 10.1101/2021.11.10.468009

**Authors:** Robert I. Horne, Ewa Andrzejewska, Parvez Alam, Z. Faidon Brotzakis, Ankit Srivastava, Alice Aubert, Magdalena Nowinska, Rebecca C. Gregory, Roxine Staats, Andrea Possenti, Sean Chia, Pietro Sormanni, Bernardino Ghetti, Byron Caughey, Tuomas P. J. Knowles, Michele Vendruscolo

## Abstract

Machine learning methods hold the promise to reduce the costs and the failure rates of conventional drug discovery pipelines. This issue is especially pressing for neurodegenerative diseases, where the development of disease-modifying drugs has been particularly challenging. To address this problem, we describe here a machine learning approach to identify small molecule inhibitors of α-synuclein aggregation, a process implicated in Parkinson’s disease and other synucleinopathies. Because the proliferation of α-synuclein aggregates takes place through autocatalytic secondary nucleation, we aim to identify compounds that bind the catalytic sites on the surface of the aggregates. To achieve this goal, we use structure-based machine learning in an iterative manner to first identify and then progressively optimize secondary nucleation inhibitors. Our results demonstrate that this approach leads to the facile identification of compounds two orders of magnitude more potent than previously reported ones.

## Introduction

Parkinson’s disease is the most common neurodegenerative movement disorder, affecting 2–3% of the population over 65 years of age^1–5^. The aggregation of α-synuclein (αS) has been associated with the initial neurodegenerative processes underlying this disease, in which the pathological accumulation of misfolded proteins results in neuronal toxicity beginning in the substantia nigra^1,2,4,6^. Since αS aggregates have been shown to exhibit various mechanisms of cellular toxicity^7,8^, major efforts are being invested into identifying compounds that can inhibit αS aggregation mechanisms^9–12^. This is a particularly pressing need given the lack of disease-modifying therapies currently available to PD patients^13–15^. With the recent approval by the FDA of the first two disease-modifying drugs for Alzheimer’s disease, aducanumab^16^ and lecanemab^17^, approaches based on blocking secondary nucleation appear to be promising^18^.

Computational methods could be expected to reduce the time and cost of traditional drug discovery pipelines^19–21^. In this area, machine learning is rapidly emerging as a powerful drug discovery strategy^22^. To explore the potential of this strategy in drug discovery programs for Parkinson’s disease and other synucleinopathies, we describe here a machine learning approach to explore the chemical space to identify compounds that inhibit the aggregation of αS. Our starting point is an approach that combines docking simulations with in vitro screening, which was recently employed to identify a set of compounds that bind to the fibril structures of αS, and prevent the autocatalytic proliferation of αS fibrils as a result^23^. Here, we used this initial set of compounds as input for a structure-based machine learning approach to identify chemical matter that is both efficacious and represents a significant departure from the parent structures, providing compounds that conventional similarity searches would have failed to efficiently identify.

This approach is based on the lessons learned using chemical kinetics about the importance of secondary nucleation in αS aggregation^24–26^. Because of the autocatalytic nature of this process, structure-based methods could be expected to effectively target the catalytic sites on the surface of αS aggregates^23^. As we show here, the implementation of this idea within an iterative machine learning procedure leads to the identification and optimisation of compounds with high hit rates and great potency.

## Results

### Components of the machine learning method

The machine learning approach used here consists of 3 main components^27^: (1) the experimental data, i.e. a readout of the potency of the compounds in an aggregation assay, (2) the variational autoencoder required to represent the compounds as latent vectors, and (3) a model for training and prediction using these vectors and the assay readouts.

For component 1, we used a chemical kinetics assay^9,28,29^ that provided both the initial data for the model training and the data that were iteratively fed back into the model at each cycle of testing and prediction. This assay identifies the top compounds that inhibit the surface-catalysed secondary nucleation step in the aggregation of αS.

For component 2, we used a junction tree variational autoencoder^30^, pre-trained on a set of 250,000 molecules^31^ enabling accurate representation of a diverse population of molecular structures. Using this approach, SMILES strings were standardised using MolVS^32^ and converted into latent vector representations.

For component 3, we used a random forest regressor (RFR) with a Gaussian process regressor (GPR) fitted to the residuals^33,34^ of the RFR, with both regressors using the latent vectors as training features. The RFR provided the highest performance compared to other combinations of multi-layer perceptrons (MLPs), GPRs and linear regressors (LRs) in terms of R^2^ score, mean absolute error (MAE) and root mean square error (RMSE). Performance and parameters are shown in **Figure S1** and **Table S1**, respectively. Combining the RFR and GPR provided only a marginal improvement in the metrics of the RFR alone, but crucially enabled leveraging of the associated uncertainty measure of the GPR when ranking molecules during acquirement prioritisation^27^. Tuning the weighting applied to this uncertainty measure allowed a ranking based on both the predicted potency of the molecules and the uncertainty of that prediction. Component 3 was then trained on the 161 initial experimental data points (see below). The best molecules predicted by the model were then tested in the same assay and the results fed back into the model in an iterative fashion (∼55-65 new molecules tested at each iteration). The molecules used at each stage of the project are illustrated in **Figure S2**, together with the structures of the most potent hits at each stage. An overview of the pipeline is shown in **Figure 1**.

**Figure 1.**
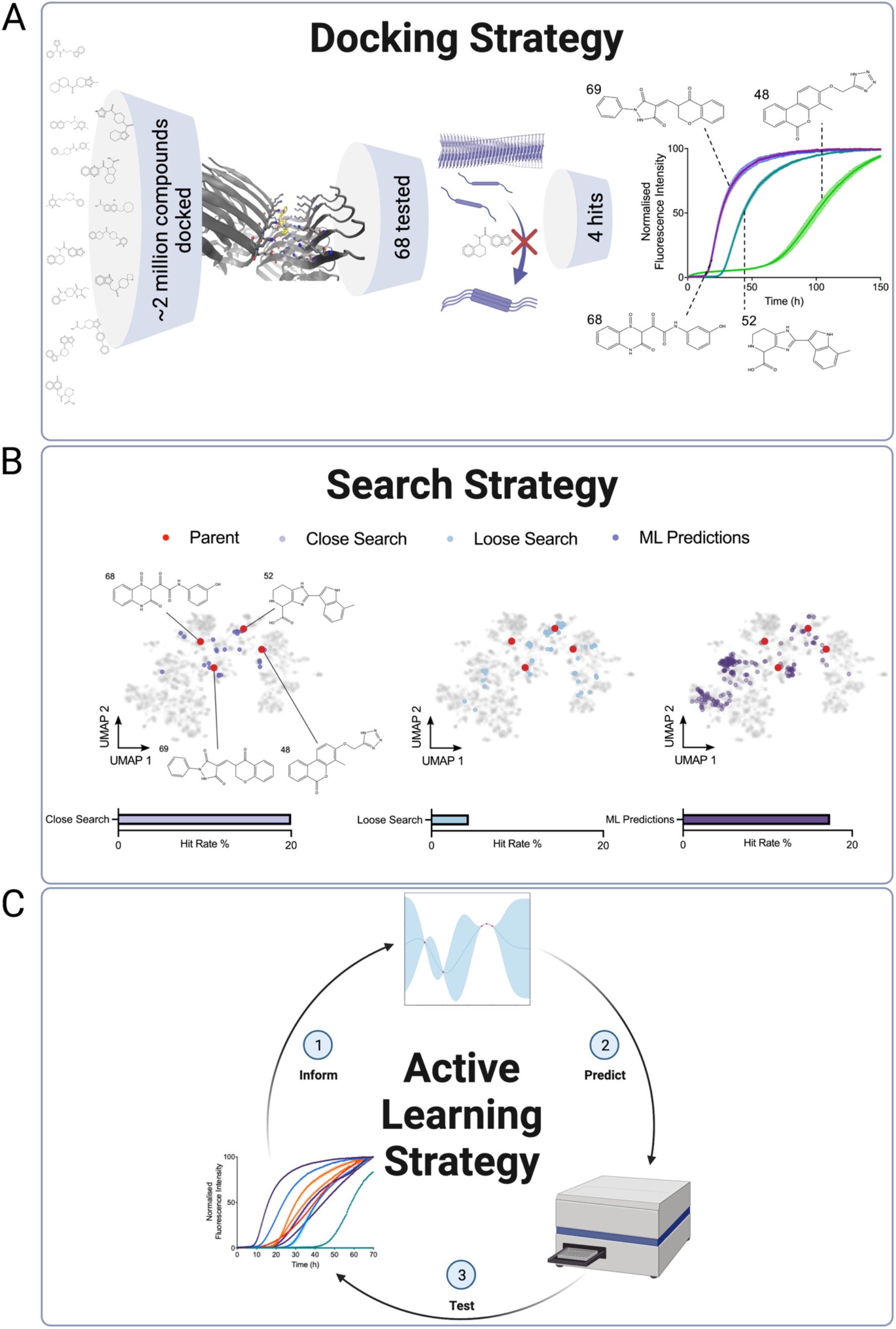
Illustration of the 3 stages of exploration of the chemical space described in this work. **(A)** From 68 molecules predicted to have good binding via docking simulations, we initially identified 4 active molecules (the ‘docking set’) by experimental testing^23^. These 4 molecules increase the *t_1/2_* of αS aggregation. **(B)** We then performed a close Tanimoto similarity search around the 4 parent compounds in chemical space. We selected molecules with Tanimoto similarity cut off > 0.5 (the ‘close similarity docking set’) followed by a loose similarity search with Tanimoto similarity cut off > 0.4 (the ‘loose similarity docking set’). A machine learning method was then applied using the observed data to predict hits from a compound library derived from the ZINC database with Tanimoto similarity > 0.3 to the parent structures (the ‘test set’). **(C)** Successive iterations of prediction and experimental testing yielded higher hit rates, and molecules with higher potency on average than those identified in the previous similarity searches. Validation experiments were also carried out on the hits identified.

### Initial set of small molecules

The initial set of molecules was identified via docking simulations to αS fibrils (see SI), followed by similarity searches around molecules that performed well in the chemical kinetics assay to identify further candidates^23^. The docking screening was carried out using the consensus strong binders predicted by AutoDock Vina^35^ and Openeye’s FRED^36–38^ software.

The first round of in vitro experiments was performed using 68 lead compounds identified in an in silico structure-based docking study carried out previously^23^. In that study, the binding site encompassing residues His50−Lys58 and Thr72−Val77 was selected due to its propensity to form a pocket according to the Fpocket software^37^ (**Figure S3A**), and its mid to low solubility according to CamSol^41^ (**Figure S3B**). Additionally, His50 is predicted to be protonated below the pH value (5.8) at which αS secondary nucleation more readily occurs^42^, which may be significant for initial interactions. 79 lead molecules were identified by docking of 2 million molecules with optimal CNS-MPO^43^ properties using Autodock Vina to target the selected binding pocket (**Figure 1A**). To increase the confidence of the calculations, the top-scoring 100,000 small molecules were selected and docked against the same αS binding site, using FRED^36^. The top-scoring, common 10,000 compounds in both docking protocols were selected and clustered using Tanimoto clustering^44^ with a similarity cut off of 0.75, leading to a list of 79 clusters. A value of 1 for the Tanimoto similarity implies complete 2D homology between 2 structures while values closer to 0 imply little to no structural similarity. 68 of the corresponding centroids of each cluster were then obtained and experimentally tested.

Subsequent experiments to test these predicted binders in aggregation assays identified 4 active compounds^23^ labelled molecule 48, 52, 68 and 69, referred to as the ‘docking set’, (**Figure 1A**). Here, using the Tanimoto similarity metric between Morgan Fingerprint representations (radius = 2, nbits = 2048) of the molecules, 2 similarity searches were then carried out using these 4 structures as starting points (**Figure 1B**). Different Tanimoto similarity thresholds were used to specify molecule subsets for testing, from initial analogue searches to the creation of a library to screen from. As such a similarity value >0.5 was used for closely related molecules, >0.4 for loosely related molecules and >0.3 for very loosely related molecules. While this use of a structurally related screening library constrains the models ability to generalise, the lack of diversity in terms of hits also makes it unlikely for the model to perform well in chemical space significantly divergent from this region. We are thus carrying out an exploitation strategy here. We remove the need for a curated screening library in a parallel work by utilising generative modelling and reinforcement learning^45^, allowing for both exploitation and exploration strategies. A selection of closely related molecules (Tanimoto similarity > 0.5) to the parent compounds (referred to as the ‘close similarity docking set’, **Figure 1B** and **Figure S2B**) was tested in the aggregation assay. The hit selection was made according to a cut off corresponding to a normalised half-time of the aggregation (*t_1/2_*) of 2 times that of the negative control. This yielded 5 new hits from 25 new molecules (**Figure S2B**), 1 derived from molecule 48, 3 from molecule 52 and 1 from molecule 69.

This step was then followed by a larger selection of compounds with a looser cut-off of structural similarity (Tanimoto similarity > 0.4) to the parent compounds (referred to as the ‘loose similarity docking set’, **Figure 1B**). Although new hits featured amongst this set, the hit rate was low (4%), and both molecules 48 and 52, which had initially appeared the most promising of the parent structures, yielded poor results. From the 29 molecules related to molecule 48 in the loose similarity docking set, none were hits, while from the 24 molecules related to molecule 52, only 2 were hits. The functional range of molecules 48 and 52 appeared narrowly limited around the chemical space of the parent structures. Molecule 69 yielded 1 hit from 16 molecules. Overall, the hit rate from the loose similarity docking set was less than a quarter of that of the close similarity docking set and involved testing 3 times as many compounds.

These results suggest that it would be challenging to further explore the chemical space using conventional structure-activity relationship (SAR) techniques without significant attrition, since the hit rate worsened as the similarity constraint to the hits was loosened. To overcome this problem, the compounds resulting from these experiments were then used as input for a machine learning method for an iterative exploration of the chemical space (**Figure 1C**). The similarity searches removed the most obvious targets of the machine learning approach, but also increased the size of the dataset available for training. The training set, however, remained small by typical machine learning standards, consisting of 161 molecules. Since training sets of this size are common in early-stage research, a further aim of this work was to demonstrate that machine learning can be used effectively even in such data sparse scenarios.

### Iterative application of the machine learning approach

One of the issues with applying machine learning to a data sparse scenario is that predictions are likely to be overconfident. While this problem can be addressed to an extent by utilising Gaussian processes, a complementary strategy is to restrict the search area to a region of chemical space that is more likely to yield successful results. To this end, a structural similarity search of the 4 hit molecules in the docking set was carried out on the ‘clean’ and ‘in stock’ subset of the ZINC database, comprising ∼6 million molecules. Any molecules showing a Tanimoto similarity value of > 0.3 to any of the 4 structures of interest was included. This low threshold for Tanimoto similarity was intended to narrow the search space but without being overly restrictive of the available chemical landscape, yielding a dataset of ∼9000 compounds which comprised the prospective ‘evaluation set’. The distribution of this evaluation set in terms of the predicting binding energies is shown in **Figure S4A**.

Different machine learning models were initially trialled against the docking scores calculated for the evaluation set as a test of the project feasibility, and these models were then tuned on the much smaller aggregation data set. The best performing set up, the RFR-GPR stacked model, was then trained on the whole aggregation data set and used to predict the top set of molecules (see Machine Learning Implementation in Supplementary Information, and **Figures S1, S5** and **S6**). For this work, the *t_1/2_* for the light seeding assay was used as the metric of potency to be used in machine learning because of its robustness. For comparison, the amplification rate is more susceptible to small fluctuations in the slope of the aggregation fluorescence trace^23^ (**Figure S7**). Molecules that achieved a *t_1/2_* 2-fold greater than that of the negative control under standard assay conditions (see Methods) were classed as hits^46^. The algorithm was run repeatedly from different random starting states and those molecules that appeared in the top 100 ranked molecules more than 50% of the time (64 molecules) were chosen for purchase (first iteration). In this first iteration, there was an inherent bias towards the structure of molecule 69 in the dataset given the relative population sizes (**Figure S2A**), but with the caveat that many of these structures were only loosely related to the parent (Tanimoto similarity < 0.4). Many of the hit molecules came from this group, suggesting chemical departures from the parent structure.

The dynamic range within the aggregation dataset in terms of potency was large, in that a majority of the molecules had no effect on aggregation, while initial docking hits exhibited relative *t_1/2_* of up to 4-5 times that of the negative control (limited by the length of the experimental run) at 25 μM. Molecules then found via machine learning produced a relative *t_1/2_* of ∼4-5 at up to 8 fold lower concentration (3.12 μM, 0.3:1 molecule:protein) than that carried out in the initial screening (25 μM, 2.5:1 molecule:protein). This compares favourably with previous molecular matter tested in a less aggressive seeded aggregation assay such as the flavone derivatives, apigenin, baicalein, scutellarein, and morin which achieved relative *t_1/2_* of 1-2 at a stoichiometry of 0.5:1 molecule:protein^9^. Anle-138b^12^ is another example of a well-characterised small molecule inhibitor, which was also taken into clinical trials, whose relative *t_1/2_* is 1.22 (**Figure 2**) at a ratio of 2.5:1 molecule:protein in the assay used in this work, which is significantly lower than any of the molecules discovered using the strategy employed here.

**Figure 2.**
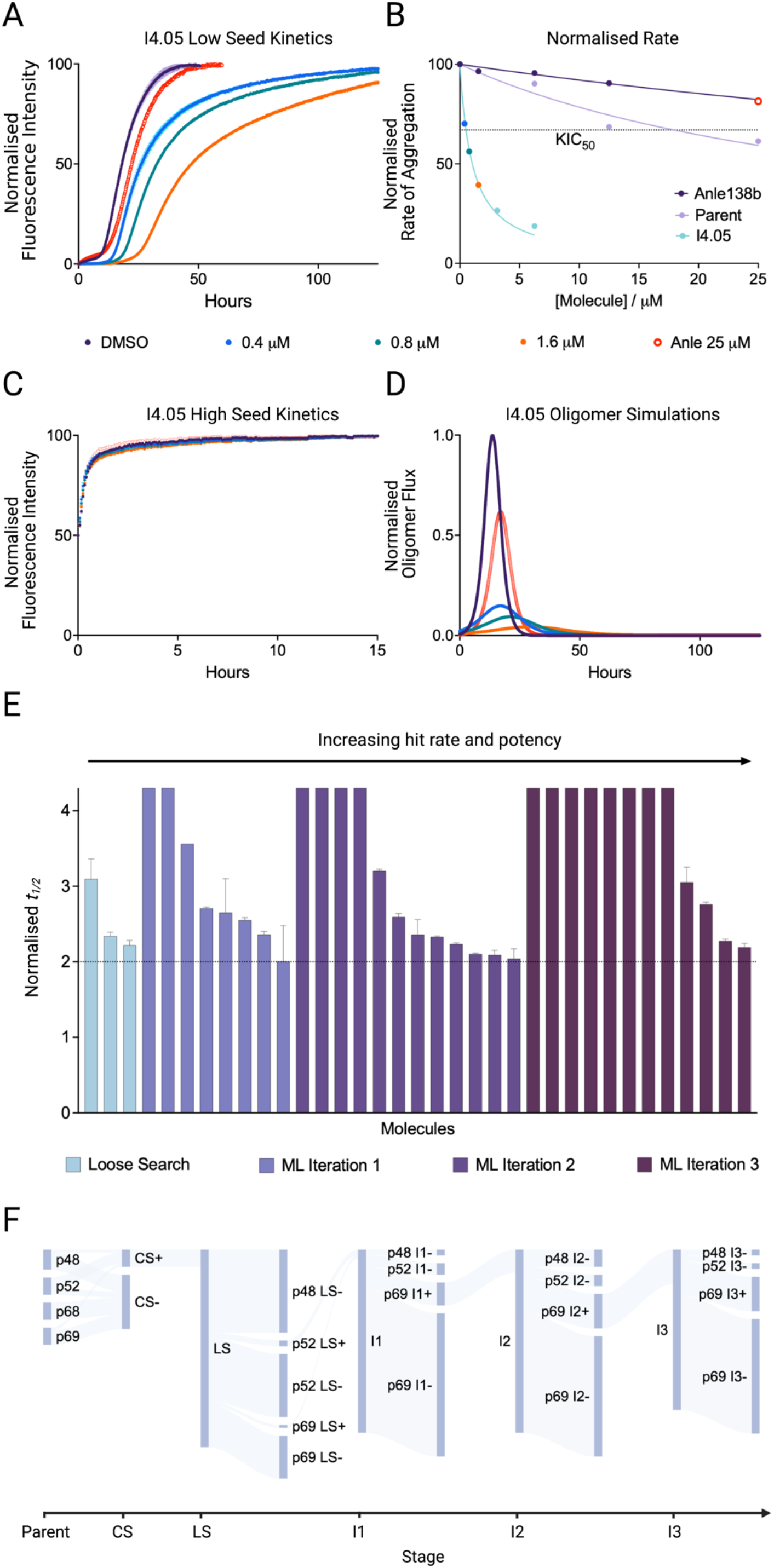
Results of the iterations of the machine learning drug discovery approach. **(A)** Kinetic traces of a 10 µM solution of αS with 25 nM seeds at pH 4.8, 37 °C in the presence of molecule or 1% DMSO in triplicate, with error bars denoting SD. During the initial screening, except for iteration 4, all molecules were screened at 2.5 molar equivalents (25 µM), and hits were then taken for further validation at lower concentrations: 0.4 µM (blue), 0.8 µM (teal), 1.6 µM (orange) with Anle-138b at 25 µM for comparison (red circles). The 1% DMSO control is shown in purple. Molecule I4.05 is shown as an example. The endpoints are normalised to the αS monomer concentration at the end of the experiment, which was detected via the Pierce™ BCA Protein Assay at *t* = 125 h. Furthermore the same experiments were carried out using AlexaFluor™ 488 labelled αS yielding similar levels of inhibition as the ThT curves. **(B)** Approximate rate of reaction (taken as 1/*t_1/2_*, normalised between 0 and 100) in the presence of 3 different molecules, Anle-138b (purple), parent structure 69 (lilac) and I4.05 (blue). The KIC_50_ of I4.05 is indicated by the intersection of the fit and the horizontal dotted line. **(C)** High seeded experiments (5 µM seeds, all other conditions match **A**) were also carried out to observe any effects on the elongation rate and enable oligomer flux calculations using the secondary nucleation rate derived from **A. (D)** Oligomer flux calculations for I4.05 vs the competitor Anle-138b using the rates derived from both **A** and **C**. **(E)** Normalised *t_1/2_* for the hits at 25 µM from the different stages: loose search, iteration 1, iteration 2 and iteration 3 (error bars denote SEM). The horizontal dotted line indicates the boundary for hit classification, which was normalised *t_1/2_* = 2. For the loose search, 69 molecules were tested, while for iterations 1, 2 and 3, the number of molecules tested was 64, 64 and 56 respectively. Note that the most potent molecules exhibited complete inhibition of aggregation over the timescale observed, so the normalised *t_1/2_* is presented as the whole duration of the experiment. **(F)** Flow of molecule hits (+) and negatives (-) in the project starting from the close search (CS), moving to the loose search (LS) and then iterations 1, 2, and 3 (I1, I2, I3). Each branch is labelled with the molecule source (e.g. p48). Attrition reached its highest point at the loose search before gradually improving with each subsequent iteration.

After the first iteration, the compound data were pooled together to extend the training set and a further 2 iterations were carried out, adding the resultant data to the training set at each iteration. This was followed by a fourth and final iteration trained on low dose (3.12 μM) data of all the previously obtained molecules. Example kinetic traces for a molecule from the fourth iteration are shown in **Figure 2A**. The molecules are labelled according to iteration number and hit identifier within that iteration. For example I4.05 is the fifth hit (05) within iteration 4 (I4). The dose-dependent potency in the aggregation assay was investigated (**Figures 2A** and **S8**) with all hit molecules exhibiting substoichiometric potency. For comparison Anle-138b is also shown. **Figure 2B** shows an approximate overall rate of aggregation at different concentrations of I4.05, Anle-138b and the parent molecule. This approximate rate was taken as 1/*t_1/2_*, and fitted to a Hill slope. A kinetic inhibitory constant (KIC_50_), the concentration of molecule at which the *t_1/2_* is increased by 50% with respect to the control, as defined previously^46^, was then derived. The KIC_50_ values for the hits were in the range of 0.5-5 μM, which compare favourably with the parent of the hit molecules (molecule 69) and Anle-138b which have extrapolated KIC_50_ values of 18.2 μM and;36.4 μM respectively. I4.05 had a KIC_50_ value of 0.52 µM with 95% confidence limits of 0.45 µM and 0.59 µM.

The elongation rate was largely unaffected in the presence of molecules at any concentration (**Figure 2C**). This was expected given the designed mechanism of action of the small molecule. It was also reassuring, since compounds that inhibit elongation may increase the population of oligomers^46^, which are considered the most damaging of the aggregate species *in vivo*^7,8^. Then, using the amplification and elongation rates derived from **Figure 2A,C**, the oligomer population over time was calculated^9^ (see Methods). These calculations are shown in **Figure 2D** for I4.05 and **Figure S8** for the rest of the hits. All hits demonstrated a dose-dependent delay and reduction of the oligomer peak. Across all metrics, I4.05 performed significantly better than Anle-138b and the parent molecule at substoichiometric ratios, as do all of the hits obtained in previous iterations (**Figures S8** and **S9**).

The aggregation data from the first 3 iterations are also shown in **Figure 2E**. Of the 64 molecules from iteration 1, 8 were strong hits, representing a hit rate of 12.5%, the second iteration showed a further increase, with 12 strong hits representing a 18.8% hit rate and the third iteration, with 12 hits, exhibited a hit rate of 21.4%. These hit rates represent an order of magnitude improvement over HTS (∼1%) and, remarkably, an overall 45% improvement over the combined similarity search hit rates, which removed the most likely hit candidates. The potency of the machine learning hits was significantly higher on average than those identified by the similarity searches (**Figure S10A**), without compromising the CNS-MPO scores (**Figure S10B**). The flow of molecules derived from each parent in terms of positives and negatives over the course of the project is illustrated in **Figure 2F**. The accumulated training data from all stages of the project for all molecules in terms of half time distribution is shown in **Figure S4B** and **S4C**.

Given that αS aggregation and toxicity has also been linked to membrane interactions^7,47^ a parallel investigation was carried out with a lipid induced aggregation assay (**Figure S11**) which was used as a validation of the molecules rather than for machine learning optimisation. The tested hit molecules also showed strong efficacy in this assay.

### Analysis of the chemical space explored by machine learning

The chemical space explored by the machine learning approach was inspected via dimensionality reduction techniques, including PCA, t-SNE^48^ and UMAP^49^ (see Methods) to investigate how the model was prioritising molecules (**Figure S12**). The relative positioning of the training points and the parents within the chemical space is shown in **Figure S13A**. The stacked RFR-GPR model assigned low uncertainty to areas of the chemical space proximal to the observed data, and the corresponding acquirement priority mirrored this when trained on the aggregation data (**Figure S13B-D**). This figure also illustrates how the uncertainty weighting could be altered during the ranking, depending on how conservative a prediction was required. A drawback to a high uncertainty penalty was that the model remained in the chemical space it was confident in, while a lower uncertainty penalty ensured reasonable confidence of hit acquirement while still exploring the chemical space.

The changes in similarity of the hits to the parent structures are shown in **Figure S14**. The similarity of the molecules to their parent structure dropped for all structures at successive stages of the investigation, reaching its lowest point at the iterations of the machine learning approach. The more potent hits mostly retained the central ring and benzene substituent of molecule 69 albeit with the addition of polar groups to the benzene ring, but featured significant alterations to the rest of the scaffold. For example, from iteration 1, I1.01 replaced the fused ring substructure of molecule 69 with a single substituted benzene ring, while I1.02 replaced it with a substituted furan ring, and subsequent iterations saw more complexity introduced. These changes were reflected in the Tanimoto similarity values, which were at the lower end of what was permitted in the evaluation set, 0.3 being the cut off. It was evident from this result that parts of the substructure were important to retain for potency, which the model did effectively while also identifying alterations in the rest of the scaffold that enhanced the potency considerably beyond that of the parent.

The observation that the QSAR model converges on the structures from two areas of the UMAP space related to structure 69 was encouraging in that it suggested the models were learning useful information and not selecting at random. While we have not tested a random set of molecules due to prohibitive resource cost, we do note that if a random selection of molecules were taken from the accumulated training data from all stages of the project, its hit rate (11%) would be lower than that of iterations 1, 2 and 3 on average. Though performance improves with additional data the QSAR performance in terms of R^2^ remains modest (**Figure S1**), but this is in part due to sparsity of training data. We would anticipate improvement if this approach could be implemented at medium scale with correspondingly more complex QSAR models, and we have an indication of this from trials of the this model set up against the docking scores of the evaluation set, where performance in terms of R^2^ score is 3 fold higher for a slightly larger dataset.

Next, an investigation was carried out to identify what structural information the latent vectors were encoding. Variational autoencoders are generally not built to ensure that their latent space dimensions are human interpretable, making this a challenge. The decoding of a variational autoencoder is also not deterministic, preventing facile analysis of the feature space based on single perturbation approaches of the input features and observing changes to decoded structures. Instead, hierarchical clustering was carried out on the latent vectors, followed by SHAP^50^ clustering for comparison (**Figure S15**). While the former differentiated groups based on large changes in any dimension, clustering based on SHAP dimensions ensured that clusters were created based only on features relevant to the prediction problem at hand. Latent space dimensions that have a large range of values had a large effect on the latent space clustering, regardless of whether these dimensions were important predictors of molecular potency. Using SHAP values, on the other hand, meant that latent space dimensions which had little effect on the model prediction were mapped to values close to zero, and therefore had a much smaller influence on the clustering. This resulted in clusters which were relevant to the prediction task. This strategy was suggested by the authors of SHAP and was recently used in the context of identifying subgroups of Covid-19 symptoms^51^.

**Figure S15** shows 2D UMAP representations of the tested molecules, with the latent vector clustering indicated by colour and the SHAP clustering indicated by shape. From the UMAP representation, we note that the SHAP clustering identified clusters more effectively than the hierarchical clustering. The SHAP values for each feature show the importance of that feature in the interpretation of potency, and this in turn could be used to identify which substructures within the molecules are relevant for potency by observing the structures that recurred in each cluster. For example, **Figure S15** shows the top dimensions of each SHAP cluster, revealing that dimension 24 at least partly encoded for the key sub-structure 3,5-pyrazolidinedione, which was present in every molecule in cluster α and a significant proportion of cluster β, while dimension 26 at least partly encoded for the key sub-structure of cluster d, a chromenone fused ring system which was present in every other molecule in the cluster. This confirmed the hypothesis previously put forward^30^ that in a junction tree variational autoencoder, the latent space encoding preserved the key features of each molecule. Molecules which were clustered together shared many molecular substructures in common.

### Measurement of binding affinity

A series of validation experiments were carried out on the most potent hits from the machine learning iterations. We first tested the binding to fibrils using surface plasmon resonance (SPR, see Methods) under different buffer conditions. The results for molecule I4.05 vs Anle-138b are shown in **Figure 3**. The proposed mechanism of action is the binding of molecules to the fibrils thereby blocking nucleation sites for further aggregation. Support for this mechanism of action comes from the observations that the molecules function at significantly substoichiometric ratios, discounting monomer interactions, and also show negligible effect on elongation. Covalent interactions can also be discounted, as no mass change is observed of the αS monomer by mass spectrometry. The large effect observed in an assay that isolates secondary nucleation as the dominant mechanism implies that the molecules are specifically affecting this step, and the substoichiometry implies that the molecules must be interacting with the fibrils which are present in nM monomer equivalents at the start of the aggregation.

**Figure 3.**
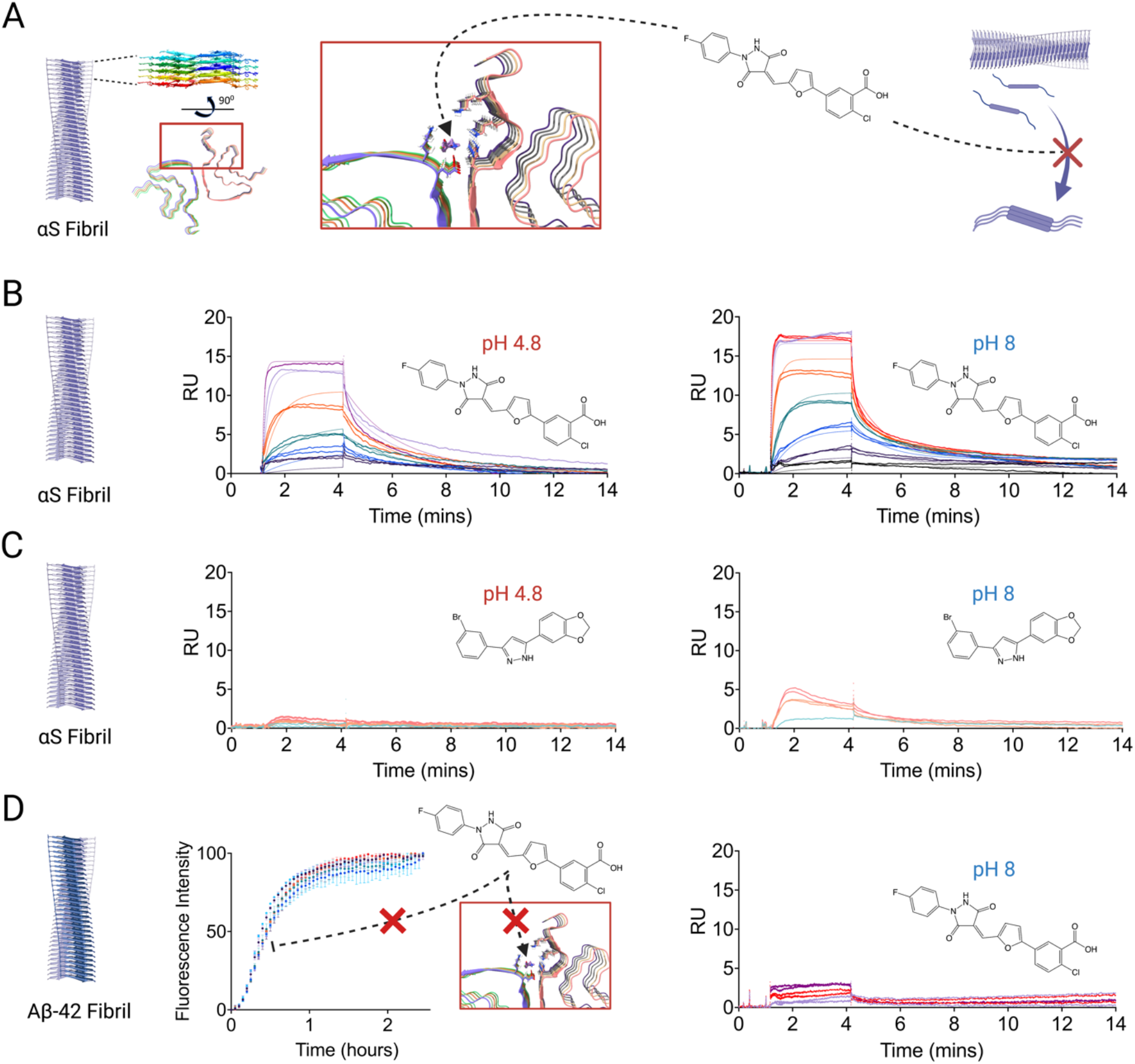
Molecule binding to αS fibrils. **(A)** A schematic representation of small molecule binding to the target binding pocket on the αS fibril, preventing secondary nucleation in the process**. (B)** SPR response curves for different concentrations of I4.05 at pH 4.8 and pH 8 binding to αS fibrils generated by a low seeded assay, with the corresponding molecular structure shown. Raw data (points) and the corresponding fits (solid lines) for each molecule concentration are shown: 1.1 nM (black), 3.3 nM (purple), 11 nM (blue), 33 nM (teal), 111 nM (orange), 333 nM (red), 500 nM (magenta) and 1.1 μM (purple). Concentrations were repeated in duplicate in a pyramidal arrangement. The αS fibrils were immobilised at a concentration of 2000 pg / mm^2^ on a CM5 Cytivia chip. The fits correspond to a 1:1 kinetic binding model, which yielded a K_D_ of 68 nM (k_a_ = 1.936 ± 0.007 10^5^ M^-1^s^-1^, k_d_ = 1.315 ± 0.003 10^-2^ s^-1^) at pH 4.8 and 13 nM at pH 8 (k_a_ = 5.879 ± 0.024 10^5^ M^-1^s^-1^, k_d_ = 0.781 ± 0.002 10^-2^ s^-1^). **(C)** SPR response curves for different concentrations of Anle-138b at pH 4.8 and pH 8 binding to αS fibrils generated by a low seeded assay, with the corresponding molecular structure shown. Raw data (points) for each molecule concentration are shown: 1.1 μM (purple), 3.3 μM (light orange), 5 μM (light red). Accurate fits at pH 4.8 could not be obtained given the low dose response, but at pH 8 a 1:1 kinetic binding model yielded an approximate K_D_ of 8.1 μM (k_a_ = 0.0359 ± 0.0005 10^5^ M^-1^s^-1^, k_d_ = 2.90 ± 0.02 10^-2^ s^-1^). **(D)** Seeded kinetics (20 nM seed) and SPR response curves for 2 μM Aß42 in the presence of 1% DMSO or different concentrations of I4.05 (colour scheme as above). I4.05 is unable to effectively inhibit Aß42 secondary nucleation or bind to Aß42 fibrils (approximate K_D_ = 2.5 μM). The Aß42 fibrils were immobilised at a concentration of 2000 pg / mm^2^ on a CM5 Cytivia chip.

Proof of binding and evidence for this potential mechanism are shown by SPR in **Figure 3**. **Figure 3A** shows a schematic representation of molecule binding to the binding pocket targeted during the initial docking simulation. **Figure 3B** shows SPR response curves for a concentration range between 0.3 nM and 1.1 μM of I4.05, while **Figure 3C** shows the same experiment utilising Anle-138b from 1.1 μM to 5 μM. The binding was tested under the conditions of the light seeded assay, pH 4.8, and also at pH 8, allowing direct comparison to the seeding assay conditions of Aß42, which were tested as a control in **Figure 3D**. αS is highly charged at neutral pH and has a PI of 4.7^52^. It therefore requires a pH in this region to render the protein uncharged in order to aggregate on an experimentally accessible timescale under quiescent conditions, whereas Aß42 is highly aggregation prone and requires high pH to prevent it aggregating too rapidly^46^. At both pH values, I4.05 exhibited binding to αS fibrils, with kinetic fits giving K_D_ values of 68 nM at the lower pH and 13 nM at the higher pH. The data for Anle-138b showed no response for pH 4.8 and so no K_D_ could be obtained, while at pH 8 an approximate K_D_ of 8.1 µM was obtained. It is evident that the 2 orders of magnitude improvement in KIC_50_ of I4.05 compared to Anle-138b was matched by a similar degree of improvement in terms of binding efficacy. **Figure 3D** shows that I4.05 has no effect on the seeded aggregation of Aß42, nor does it bind effectively to Aß42 fibrils, which suggests that this molecule is not a promiscuous aggregation inhibitor between different amyloidogenic proteins.

### Inhibition of aggregation using brain-derived seeds

While this result was encouraging, with the recent determination of the pathological αS fibril structure^39^, it became clear that the recombinant *in vitro* fibril structure we had employed for computational and experimental work was different to that found in the brains of Parkinson’s disease patients. To test whether these molecules might work against patient-derived fibrils, these molecules were tested in an RT-QuIC assay (**Figure 4**) that employs brain samples from patients suffering with Dementia with Lewy Bodies (DLB). The fibril structure found in DLB was found to match that found in Parkinson’s disease^39^.

**Figure 4.**
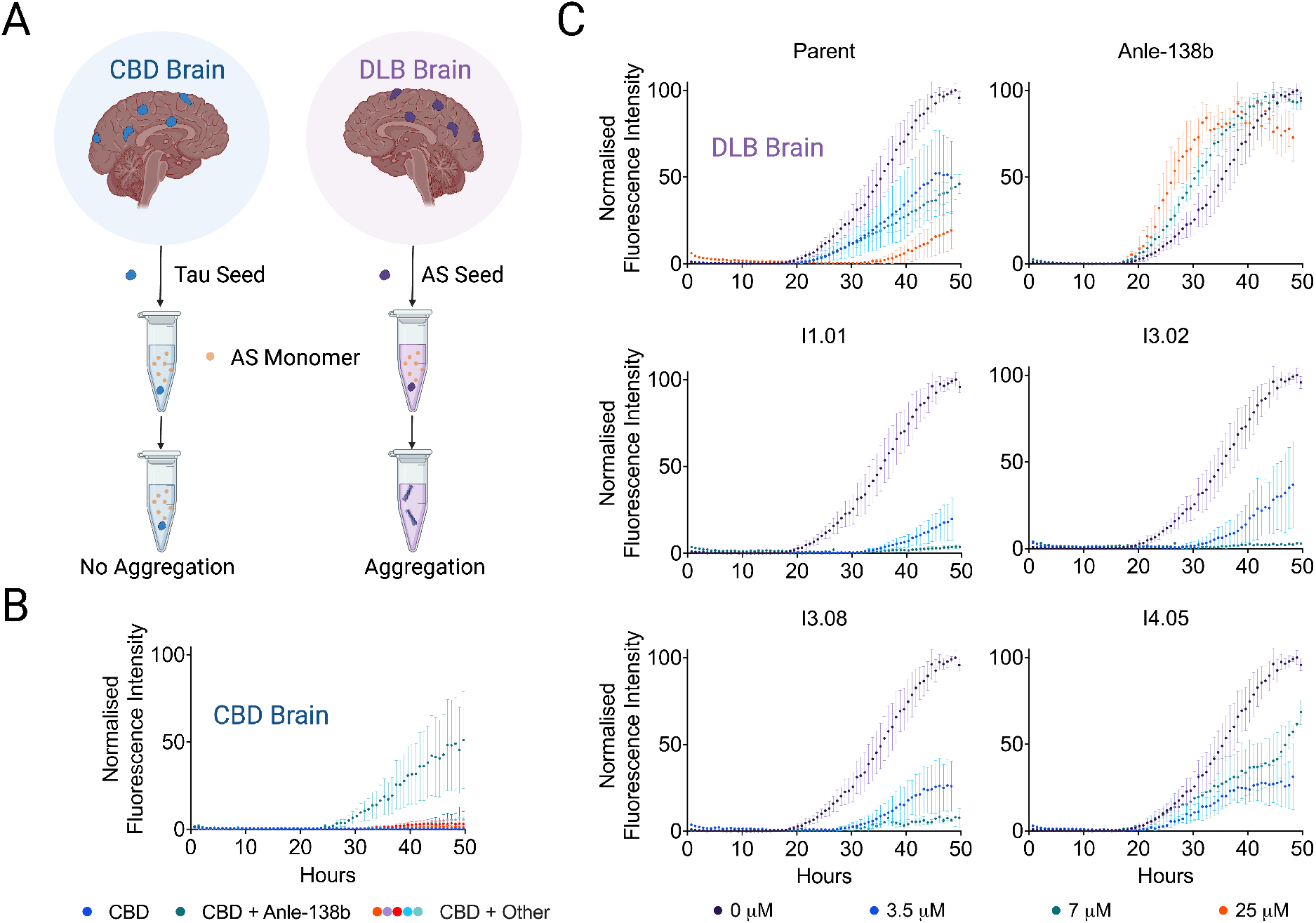
RT-QuIC brain seeding assay. **(A)** Schematic representation of the RT-QuIC assay, aggregates derived from the brain tissue of patients suffering with dementia with Lewy bodies (DLB) were used to induce αS aggregation. Samples from brains of patients with corticobasal degeneration (CBD) were used as a negative control. **(B)** Kinetic traces of a 7 µM solution of αS in the presence of CBD seeds (pH 8, 42°C, shaking at 400 rpm with 1 min intervals, in quadruplicate, error bars denote SD). CBD samples were 1% DMSO (blue), 7 µM Anle-138b (teal), parent (orange), I1.01 (purple), I3.02 (red), I3.08 (turquoise) and I4.05 (light blue). Anle-138b, in teal, induces aggregation under this condition. **(C)** Kinetic traces of a 7 µM solution of αS in the presence of DLB seeds. The DLB samples were 1% DMSO (purple), 3.5 µM molecule (blue), 7 µM molecule (teal) and 25 µM molecule (orange). Anle-138b again appears to accelerate rather than inhibit aggregation.

The RT-QuIC assay was initially introduced as a diagnostic assay^53,54^, showing distinct aggregation curves in the presence of brain material derived from different pathologies^55^. In this case, we use it to test the ability for these molecules to slow the aggregation of αS induced by DLB brain material. As a negative control, samples from patients with a tauopathy (corticobasal degeneration, CBD) were also used, as these did not induce αS aggregation as no αS seeds were present (**Figure 4A, B**). The conditions are different to those initially screened, as this assay was carried out at pH 8 and utilised shaking to induce aggregation. This is a more challenging paradigm for the molecules to function in as multiple aggregation processes occur in tandem^42^. In addition to secondary nucleation from the fibril surfaces, fragmentation of the fibrils induced via shaking results in more fibril ends for elongation, which in turn provides more fibril surface for secondary nucleation.

Despite these challenges, and the different fibril structure present, the molecules still function well in inhibiting aggregation, and still at substoichiometric ratios (**Figure 4C**). There is again a clear improvement for the hits over Anle-138b, which in fact appears to accelerate aggregation in this example, and the parent molecule, although the ranking of the hits in terms of efficacy is altered compared to the screening assay. To understand these results we note that there is a similarity in the binding pockets in the structures 6CU7 (recombinant) and 8A9L (brain derived) (**Figure S16**). We currently do not know whether or not this similarity is serendipitous, but binding pockets with similar features can also be observed via cryo-EM in the MSA-I and MSA-II fibril folds as well as the Lewy fold, with an unresolved species bound within the pocket^39^.

To account for differences in brain samples and also investigate potential efficacy against MSA derived brain material, we tested a single concentration of the same selection of molecules against 3 neuropathologically confirmed MSA brain samples (**Figure S17A, C**) and 2 further DLB brain samples (**Figure S17A, D**). As a further negative control, a sample with no seed was tested, to determine the degree of spontaneous nucleation in the absence of brain material (**Figure S17B**). Aggregation in the negative control is effectively inhibited by all the ML molecules, given αS is likely to assume the 6CU7 polymorph in this condition, and not by Anle-138b which accelerates aggregation. It should be noted however that the CBD samples are the better negative control for RT-QuIC, as all brain samples contain traces of cell matrix components that may sequester αS and reduce its aggregation. The unseeded sample begins aggregation at ∼40-50 h whereas CBD samples do not exhibit significant aggregation over a span of 80 h (**Figure S17E**). Fibrils present in DLB and MSA samples are able to counteract this effect. For the DLB and MSA samples broadly similar trends were observed to those shown in **Figure 4**. The ML molecules did appear more efficacious against MSA samples (**Figure S17C**), perhaps because the MSA pocket more closely matches that of the targeted 6CU7 polymorph (4 flanking lysines around a histidine residue) compared to the 8A9L polymorph found in PD and DLB (4 flanking lysines around a tyrosine residue) as shown in **Figure S16**. The behaviour of Anle-138b was variable as, where the ML derived molecules inhibited aggregation to some extent across all examples, Anle-138b either had no effect (unseeded and MSA samples 1 and 2) or induced (CBD sample, MSA sample 3 and DLB sample 1) or mildly inhibited aggregation (DLB samples 2 and 3). No aggregation was observed in the CBD samples over the time scale observed except for Anle-138b, which accelerated aggregation under this condition.

### Oligomer quantification by micro free-flow electrophoresis

Having observed that molecule I3.02 was the most broadly effective in the RT-QuIC assay, an investigation was carried out to directly measure the oligomeric species formed during the reaction. This was achieved using microfluidic free-flow electrophoresis (µFFE)^56^, a technique optimised using similar conditions to that used in the RT-QuIC assay, albeit at significantly higher αS concentration (100 µM). The results of this are shown in **Figure 5**. Aggregation time courses were tracked using AlexaFluor™ 488 labelled N122C rather than ThT. **Figure 5** shows a schematic of the approach, where samples were extracted from an aggregation time course, centrifuged to remove insoluble aggregates, and finally submitted to µFFE. The degree of deflection and the photon count of each particle are proportional to the size and charge of the biomolecule. The former allows the separation of monomers from oligomers and the latter gives a measure of the number and size of the oligomers at a particular time point in the presence of different inhibitors. Oligomer electrophoretic mobility (*μ*_o_) for an oligomer comprised of *n*_m_ monomer units is proportional to oligomer charge (*q*_o_) and inversely proportional to oligomer hydrodynamic radius (*r*_o_) and so can be described by^56^

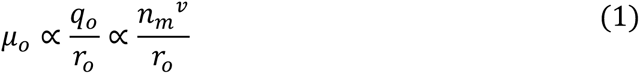

where *v* is a scaling exponent linking *q*_o_ with *n*_m_. Approximating the oligomers as spherical species yields^56^

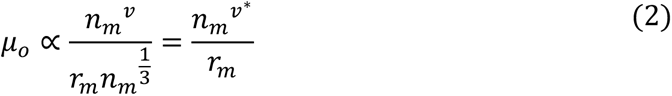

where the oligomer electrophoretic mobility is defined only in terms of the monomer number (*n*_m_) and hydrodynamic radius (*r*_m_), and the scaling exponent *v\** = *v* - 1/3. Samples were extracted at the *t_1/2_* of the negative control (1% DMSO) and the results are shown in **Figure 5**. Anle-138b dosing resulted in a smaller population of large aggregates, as may be expected from the slight acceleration in the aggregation observed in the fluorescence values, while I3.02 reduced both the size and the number of oligomers present in comparison to the DMSO control.

**Figure 5.**
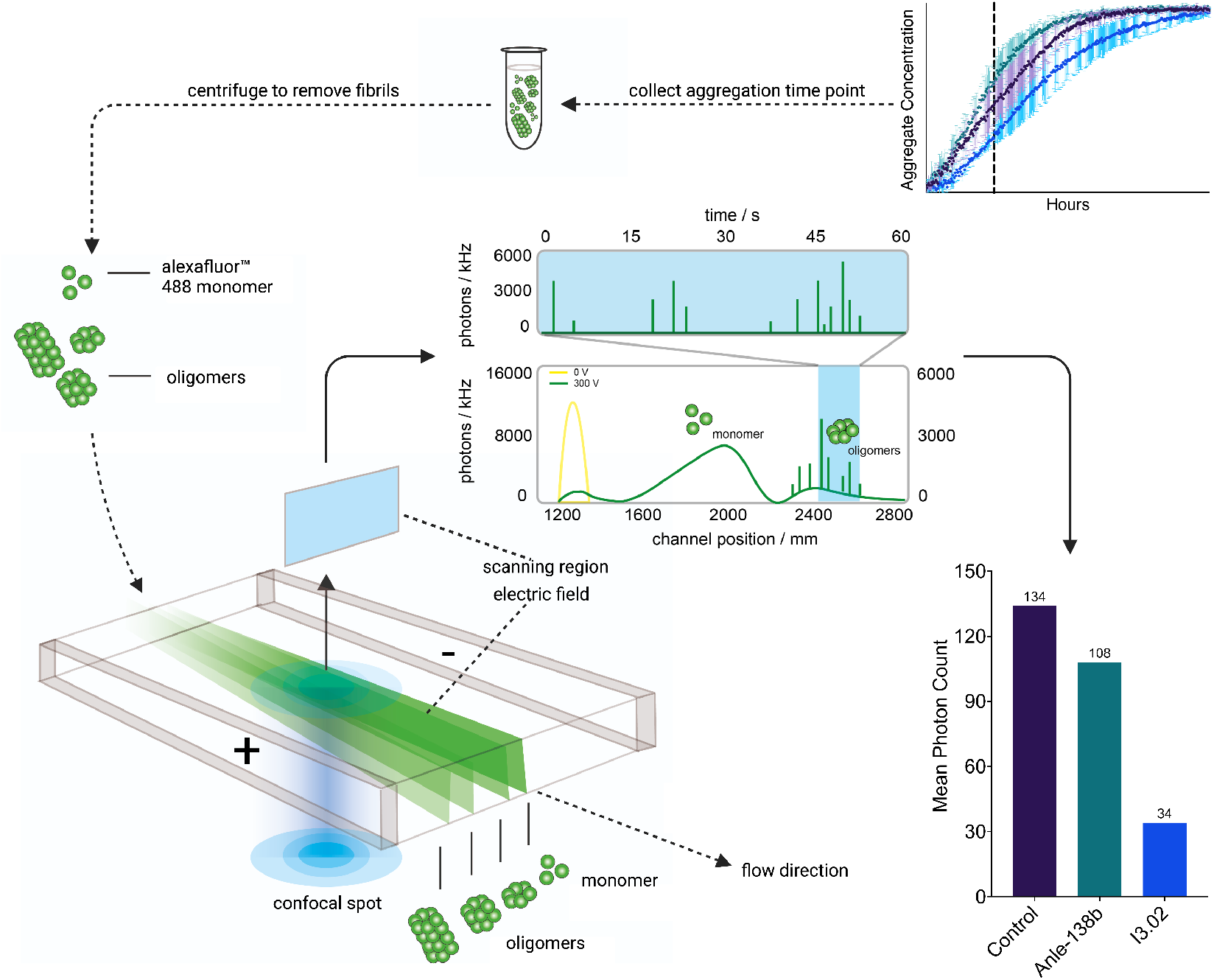
Quantification of αS oligomers using micro free-flow electrophoresis (μFFE). (Top right) αS labelled with AlexaFluor™ 488 (100 µM, pH 7.4, 37°C, cycles of 5 min shaking at 200 rpm and 1 min rest, in quadruplicate, error bars denote SD) was supplemented with 0.5 µM seed and 1% DMSO (purple) or 50 µM Anle-138b (teal) or I3.02 (blue) in 1% DMSO. Anle-138b slightly accelerates aggregation under these conditions, where fragmentation mechanisms may again play a role due to shaking, while I3.02 slows it down. Samples were extracted at 9 h from the time course of aggregation and centrifuged to remove fibrils from the mixture, leaving only αS monomers and soluble oligomeric species for analysis via μFFE. (Bottom left) Schematic representation of the μFFE approach, showing the AlexaFluor™ 488-labeled αS oligomeric mixture undergoing μFFE. The direction of fluid flow is shown by arrows. The differential deflection of the electric field allows the monomer population to be separated from the oligomer population during analysis (Middle and bottom right). Analysis of the aggregate populations detected in each sample. The number of photons emitted, proportional to particle number and size, is plotted on the y axis of the bar plot for each sample. The average number of photons emitted per particle is indicated in the inset.

The identification of inhibitors of αS aggregation based on chemical kinetics approaches has advanced to the point that specific steps in the aggregation process, including primary nucleation and secondary nucleation, can be targeted in a reproducible way^9,28,29^. The mechanism targeted in this work is the surface-catalysed secondary nucleation step, which is responsible for the autocatalytic proliferation of αS fibrils. In a recent initial report, initial hit molecules identified via docking simulations were shown to bind competitively with αS monomers along specific sites on the surface of αS fibrils^23,24,57^. Specific rate measures and other aggregation metrics were derived from these experiments allowing quantitative and reliable comparisons between molecules in terms of SAR and offering metrics to optimise structures of interest^9,46^. This has been augmented with tests against diseased brain material and detailed, experimental fibril binding and oligomer flux analyses.

## Discussion

The aim of this work was to develop a machine learning approach to drug discovery for protein aggregation diseases that could improve both the hit rate of the in vitro assays employed and provide novel chemical matter more efficiently than conventional approaches. As of the first iterations, the hit rate of the approach using initial hit compounds identified via docking simulations was an over 20-fold improvement over typical HTS hit rates (∼0-1%)^58^. These structures also represent discoveries that could not have been obtained by staying close in chemical space to the parent structure, as would have been dictated by similarity search approaches. There were ∼4000 molecules in the test set that have Tanimoto similarity values in the range of these hits, and all of these would potentially have had to be screened to locate these hits using similarity searches alone, as demonstrated by the looser similarity search approach which exhibited a comparatively poor hit rate (4%) despite more conservative structural alterations to the parent hits. The machine learning method was therefore able to supply a degree of novelty as well as an improved hit rate.

A limitation of this approach is the requirement to select molecules from a pre-existing library. To resolve this limitation generative modelling combined with reinforcement learning has been applied in a parallel project to remove the need for a library to screen from^45,59^. A second limitation is the focus on one assay metric of interest as a learning parameter. Addressing this limitation will involve future work on multi-parameter optimisation, which is a challenging area in rapid development^60–62^. Another topic of great interest in drug discovery approaches based on machine learning besides potency prediction is the prediction of pharmacokinetics and toxicity^63^. It could be possible to achieve this multi-parameter optimisation utilising multiple models in parallel and then employing a joint ranking metric, or architectures that screen for individual metrics in series, although this has primarily been demonstrated with predicted chemical properties such as clogP and QED rather than experimental results^60–62^. The molecules in this work were derived from a set that passed CNS-MPO criteria in the initial docking simulation, and so the CNS-MPO score of the whole aggregation inhibitor set is relatively favourable with most hit molecules exceeding the common cut off value of 4^43^ (**Figure S10B**).

It would have been preferable to begin this approach using seeds derived from relevant pathological brain material, but this was not possible, as neither structures nor samples for these were available at the start of this study. Nonetheless, we have demonstrated that these molecules still function against disease relevant inducers, likely because of the degree of commonality between the binding sites of the fibril polymorphs. The complete loss of function against another aggregation prone protein, Aß42, does however suggest specific functionality against αS alone.

## Conclusions

The results that we have presented illustrate a drug discovery approach that involves an iterative structure-based machine learning strategy to generate potent protein aggregation inhibitors. The resulting hits offer a significant improvement in potency over the parent and competitor molecules and represented a major structural departure from them. We anticipate that using machine learning approaches of the type described here could be of significant benefit to researchers working in the field of protein misfolding diseases, and indeed early-stage drug discovery research in general.

## Materials and Methods Compounds and chemicals

Compounds were purchased from MolPort (Riga, Latvia) or Mcule (Budapest, Hungary) and prepared in DMSO to a stock of 5 mM. All chemicals used were purchased at the highest purity available.

### Recombinant αS expression

Recombinant αS was purified based on previously described methods^25,42,64^. The plasmid pT7-7 encoding human αS was transformed into BL21 (DE3) competent cells. Following transformation, the competent cells were grown in 6L 2xYT media in the presence of ampicillin (100 μg/mL). Cells were induced with IPTG, grown overnight at 28 °C and then harvested by centrifugation in a Beckman Avanti JXN-26 centrifuge with a JLA-8.1000 rotor at 5000 rpm (Beckman Coulter, Fullerton, CA). The cell pellet was resuspended in 10 mM Tris, pH 8.0, 1 mM EDTA, 1 mM PMSF and lysed by sonication. The cell suspension was boiled for 20 min at 85 °C and centrifuged at 18,000 rpm with a JA-25.5 rotor (Beckman Coulter). Streptomycin sulfate was added to the supernatant to a final concentration of 10 mg/mL and the mixture was stirred for 15 min at 4 °C. After centrifugation at 18,000 rpm, the supernatant was taken with an addition of 0.36 g/mL ammonium sulfate. The solution was stirred for 30 min at 4 °C and centrifuged again at 18,000 rpm. The pellet was resuspended in 25 mM Tris, pH 7.7, and the suspension was dialysed overnight in the same buffer. Ion-exchange chromatography was then performed using a Q Sepharose HP column of buffer A (25 mM Tris, pH 7.7) and buffer B (25 mM Tris, pH 7.7, 1.5 M NaCl). The fractions containing αS were loaded onto a HiLoad 26/600 Superdex 75 pg Size Exclusion Chromatography column, and the protein (≈ 60 ml @ 200 µM) was eluted into the required buffer. The protein concentration was determined spectrophotometrically using ε280 = 5600 M^−1^ cm^−1^. The cysteine-containing variant (N122C) of αS was purified by the same protocol, with the addition of 3 mM DTT to all buffers.

### Labelling of αS

αS protein was fluorophore-labelled to enable visualisation by fluorescence microscopy. In order to remove DTT, cysteine variants of αS were buffer exchanged into PBS or sodium phosphate buffer by use of P10 desalting columns packed with Sephadex G25 matrix (GE Healthcare). The protein was then incubated with an excess of AlexaFluor™ 488 dye with maleimide moieties (Thermofisher Scientific) (overnight, 4 °C on a rolling system) at a molar ratio of 1:1.5 (protein-to-dye). The labelling mixture was loaded onto a Superdex 200 16/600 (GE Healthcare) and eluted in PBS buffer at 20 °C, to separate the labelled protein from free dye. The concentration of the labelled protein was estimated by the absorbance of the fluorophores, assuming a 1:1 labelling stoichiometry (AlexaFluor™ 488: 72000 M^-1^ cm^-1^ at 495 nm).

### αS seed fibril preparation

αS fibril seeds were produced as described previously^25,42^. Samples of αS (700 µM) were incubated in 20 mM phosphate buffer (pH 6.5) for 72 h at 40 °C and stirred at 1,500 rpm with a Teflon bar on an RCT Basic Heat Plate (IKA, Staufen, Germany). Fibrils were then diluted to 200 µM, aliquoted and flash frozen in liquid N_2_, and finally stored at −80 °C. For the use of kinetic experiments, the 200 µM fibril stock was thawed, and sonicated for 15 s using a tip sonicator (Bandelin, Sonopuls HD 2070, Berlin, Germany), using 10% maximum power and a 50% cycle.

### Measurement of αS aggregation kinetics

αS was injected into a Superdex 75 10/300 GL column (GE Healthcare) at a flow rate of 0.5 mL/min and eluted in 20 mM sodium phosphate buffer (pH 4.8) supplemented with 1 mM EDTA. The obtained monomer was diluted in buffer to a desired concentration and supplemented with 50 µM ThT and preformed αS fibril seeds. The molecules (or DMSO alone) were then added at the desired concentration to a final DMSO concentration of 1% (v/v). Samples were prepared in low-binding Eppendorf tubes, and then pipetted into a 96-well half-area, black/clear flat bottom polystyrene NBS microplate (Corning 3881), 150 µL per well. The assay was then initiated by placing the microplate at 37 °C under quiescent conditions in a plate reader (FLUOstar Omega, BMG Labtech, Aylesbury, UK). The ThT fluorescence was measured through the bottom of the plate with a 440 nm excitation filter and a 480 nm emission filter. After centrifugation at 5000 rpm to remove aggregates the monomer concentration was measured via the Pierce™ BCA Protein Assay Kit according to the manufacturer’s protocol.

For the lipid induced assay, small unilamellar vesicles (SUVs) containing 1,2-dimyristoyl-sn-glycero-3-phospho-L-serine (DMPS), Avanti Polar Lipids Inc., Alabaster, AL, USA), were prepared from chloroform solutions of the lipids as described previously^64^. Briefly, the lipid mixture was evaporated under a stream of nitrogen gas and then dried thoroughly under vacuum to yield a thin lipid film. The dried thin film was re-hydrated by adding aqueous buffer (20 mM sodium phosphate, pH 6.5, 1 mM EDTA) at a concentration of 1 mM and heating to 40 °C for 2 h while stirring at 1,500 rpm with a Teflon bar on an RCT Basic Heat Plate (IKA, Staufen, Germany). SUVs were obtained using several cycles of freeze-thawing followed by extrusion through membranes with 200 nm diameter pores (Avanti Polar Lipids, Inc). αS was prepared as above. Kinetic conditions were 20 µM αS, 100 µM DMPS, 50 µM ThT, 30 °C, all other conditions remained the same as above.

Transmission electron microscopy (TEM) imaging of the fibrils produced at the end of the light seeded aggregation reaction (**Figure S16**), was used to verify fibrils were produced

### Determination of the αS elongation rate constant

In the presence of high concentrations of seeds (≈ µM), the aggregation of αS is dominated by the elongation of the added seeds^25,42^. Under these conditions where other microscopic processes are negligible, the aggregation kinetics for αS can be described by^9,23,25^

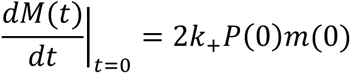

where *M(t)* is the fibril mass concentration at time *t*, *P(0)* is the initial number of fibrils, *m(0)* is the initial monomer concentration, and *k_+_* is the rate of fibril elongation. In this case, by fitting a line to the early time points of the aggregation reaction as observed by ThT kinetics, 2*k_+_P(0)m(0)* can be calculated for αS in the absence and presence of the compounds. Subsequently, the elongation rate in the presence of compounds is expressed as a normalised reduction as compared to the elongation rate in the absence of compounds (1% DMSO).

### Determination of the αS amplification rate constant

In the presence of low concentrations of seeds (∼ nM), the fibril mass fraction, *M(t)*, over time was described using a generalised logistic function to the normalised aggregation data^9,65^

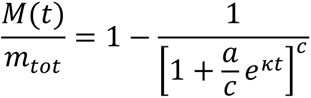

where *m_tot_* denotes the total concentration of αS monomers. The parameters a and c are defined as

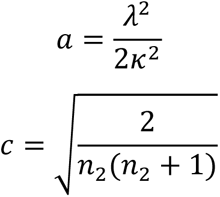

The parameters *λ* and *κ* represent combinations for the effective rate constants for primary and secondary nucleation, respectively, and are defined as^65^

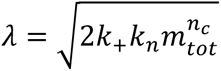

and

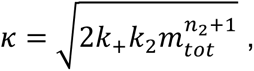

where *k_n_* and *k_2_* denote the rate constants for primary and secondary nucleation, respectively, and *n_c_* and *n_2_* denote the reaction orders of primary and secondary nucleation, respectively. In this case, *n_c_* was fixed at 0.3 for the fitting of all data (corresponding to a reaction order of *n*_2_ = 4), and *k_2_*, the amplification rate, is expressed as a normalised reduction for αS in the presence of the compounds as compared to in its absence (1% DMSO).

### Determination of the αS oligomer flux over time

The theoretical prediction of the reactive flux towards oligomers over time was calculated as^9,65^

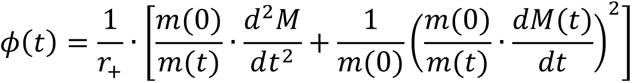

where *r_+_ = 2k_+_m(0)* is the apparent elongation rate constant extracted as described earlier, and *m(0)* refers to the total concentration of monomers at the start of the reaction.

### Recombinant Aß42 expression

The recombinant Aß42 peptide (MDAEFRHDSGY EVHHQKLVFF AEDVGSNKGA IIGLMVGGVV IA), here called Aß42, was expressed in the E. coli BL21 Gold (DE3) strain (Stratagene, CA, U.S.A.) and purified as described previously. Briefly, the purification procedure involved sonication of E. coli cells, dissolution of inclusion bodies in 8 M urea, and ion exchange in batch mode on diethylaminoethyl cellulose resin followed by lyophylisation. The lyophilised fractions were further purified using Superdex 75 HR 26/60 column (GE Healthcare, Buckinghamshire, U.K.) and eluates were analysed using SDS-PAGE for the presence of the desired peptide product. The fractions containing the recombinant peptide were combined, frozen using liquid nitrogen, and lyophilised again.

### Aß42 aggregation kinetics and fibril preparation

Solutions of monomeric Aß42 were prepared by dissolving the lyophilized Aß42 peptide in 6 M guanidinium hydrocholoride (GuHCl). Monomeric forms were purified from potential oligomeric species and salt using a Superdex 75 10/300 GL column (GE Healthcare) at a flowrate of 0.5 mL/min, and were eluted in 20 mM sodium phosphate buffer, pH 8 supplemented with 200 µM EDTA and 0.02% NaN3. The centre of the peak was collected and the peptide concentration was determined from the absorbance of the integrated peak area using ε280 = 1490 l mol^-1^ cm^-1^. The obtained monomer was diluted with buffer to the desired concentration and supplemented with 20 μM thioflavin T (ThT) from a 2 mM stock. Each sample was then pipetted into multiple wells of a 96-well half-area, low-binding, clear bottom and PEG coated plate (Corning 3881), 80 µL per well, in the absence and the presence of different molar-equivalents of small molecules (1% DMSO). Assays were initiated by placing the 96-well plate at 37 °C under quiescent conditions in a plate reader (Fluostar Omega, Fluostar Optima or Fluostar Galaxy, BMGLabtech, Offenburg, Germany). The ThT fluorescence was measured through the bottom of the plate using a 440 nm excitation filter and a 480 nm emission filter. Fibrils were extracted directly from wells and used on the day for SPR experiments.

### Machine learning

Code Availability:

Full code can be found on the GitHub repository: https://github.com/rohorne07/Iterate

#### Junction tree neural network variational autoencoder

The autoencoder^30^ was pretrained on a library of 250,000 compounds^31^, and was implemented as described previously^30^ using a pip installable version (https://github.com/LiamWilbraham/jtnnencoder). Any molecules that contained substructures the autoencoder could not represent (i.e. that fell outside the substructure vocabulary of the pretrained model) were excluded.

#### Prediction module

All coding was carried out in Python 3. Scikit-learn^66^ implementations of the Gaussian process regressor (GPR), random forest regressor (RFR), linear regressor (LR) and multi-layer perceptron (MLP) methods were tested in various combinations, and the results are shown in the supplementary section. For data handling, calculations and graph visualisation the following software and packages were used: pandas^67^, seaborn^68^, matplotlib^69^, numpy^70^, scipy^71^, umap-learn^49^, Multicore-TSNE^48^ and GraphPad Prism 9.1.2. Cross validation and benchmarking were also carried out for each model using scikit-learn built in functions and is described in the results section.

#### SHAP and latent space clustering

To compute the SHAP values, we used the SHAP python library^50^. The pretrained random-forest model was loaded, and a SHAP explainer object was created and provided with the latent representation for the top 100 highest predicted molecules. This allowed for the identification of dimensions important to the prediction of high potency molecules. The full testing set derived from the ZINC dataset was also used in order to differentiate between dimensions important to distinguish high potency molecules from low potency molecules versus dimensions important to distinguish high potency molecules between themselves. This resulted in a global interpretation of the model, encompassing all data points passed to the explainer object. The resultant plots were generated using SHAP built-in plot functions. The sklearn library hierarchical clustering method was used to cluster latent vectors for comparison, with initial cluster number set to 7^72^.

### Surface plasmon resonance

All work was carried out using Biacore T200 at 25 °C. CM5 chips were activated by flowing 0.01 M NHS, 0.4 M EDC at a flow rate of 10 µL / min for 7 minutes over 2 lanes. Preformed αS or Aß42 fibrils (derived from the endpoints of low seeded aggregation reactions) at a concentration of 1 µM in sodium acetate (10 mM, pH 4.0) were injected onto a single lane in 60s bursts at 5 µL / min until a response of 2000 units was reached. Both lanes were then deactivated using a 7-minute injection of ethanolamine (1 M, pH 8.5) at 10 µL / min, and the reference lane signal was subtracted from the active lane. Different small molecule concentrations were then flowed over both lanes in a pyramidal arrangement in duplicate with blank subtraction (association time = 3 minutes, dissociation time = 10 minutes). The running buffer was sodium phosphate (20 mM, 1 mM EDTA, variable pH) with 1% DMSO. Fitting was carried out on Biacore T200 Evaluation Software, version 3.2, using a 1:1 binding model with the RI (RU) set to a constant value of 0.

### Preparation of human brain tissue homogenates

Deidentified postmortem human brain specimens used in the RT-QuIC assay are referenced in **Table S2**. These specimens were obtained from the NIH Brain & Tissue repository-California, Human Brain & Spinal Fluid Resource Centre, VA West Los Angeles Medical Center, Los Angeles, California which is supported in part by National Institutes of Health and the US department of Veterans Affairs. Assay samples were prepared as 10% (wt/vol) brain homogenates in ice-cold phosphate-buffered saline (PBS) (pH 7.0) using 1 mm zirconia beads (BioSpec, cat#11079110z) in a Bead Mill 24 (Fisher Scientific). Subsequent dilutions of each brain homogenate (10^-1^ to 10^-5^) for testing in the RT-QuIC assay were prepared in 1X PBS (pH 7.0).

### αSyn RT-QuIC protocol

RT-QuIC assay for DLB samples were performed using the recombinant αSyn K23Q substrate purified using a 2-step chromatography protocol described previously (PMID: 29422107). For testing MSA samples, wild type αSyn (WT) recombinant substrate was purified using anion-exchange and size exclusion chromatography as described previously with minor modifications (PMID: 15939304). The WT protein expressing pET21a-αS plasmid was a gift from Michael J Fox Foundation MJFF (Addgene plasmid # 51486; http://n2t.net/addgene:51486; RRID:Addgene_51486). RT-QuIC assay was performed using black, clear bottom 96-well plates (Nalgene Nunc International) preloaded with 6 silica beads (1 mm diameter, OPS Diagnostics). Seeding was induced by addition of 2 μL of 10^-4^ (with respect to solid brain tissue) dilutions of DLB, MSA, or CBD (control) brain homogenates in quadruplicate wells containing 98 μL of the reaction buffer (40 mM phosphate buffer; pH 8.0 and 170 mM NaCl) supplemented with 6 μM (0.1 mg/ml) αSyn K23Q substrate (prefiltered through 100 kDa MWCO filter, Pall Corporation, Catalogue# OD100C34) and 10 μM ThT. After seeding, reaction plates were covered with a sealer film (Nalgene Nunc International) and incubated at 42 °C in a fluorescence plate reader (BMG FLUOstar Omega) with 1 min shake-rest cycles (400 rpm double orbital) for 50-90 h as indicated in the figures. ThT fluorescence (λ_excitation_; 450 +/− 10 nm and λ_emission_; 480 +/− 10 nm) was measured at 45 min intervals).

### Microfluidic free-flow electrophoresis

#### Microfluidic device fabrication

Devices were designed using AutoCAD software (Autodesk) and photolithographic masks printed on acetate transparencies (Micro Lithography Services). Polydimethylsiloxane (PDMS) devices were produced on SU-8 moulds fabricated via photolithographic processes as described elsewhere^75,76^ with UV exposure performed with custom-built LED-based apparatus^77^. Following development of the moulds, feature heights were verified by profilometer (Dektak, Bruker) and PDMS (Dow Corning, primer and base mixed in 1:10 ratio) applied and degassed before baking at 65 °C for 1.5 h. Devices were cut from the moulds and holes for tubing connection (0.75 mm) and electrode insertion (1.5 mm) were created with biopsy punches, the devices were cleaned by application of Scotch tape and sonication in IPA (5 min). After oven drying, devices were bonded to glass slides using an oxygen plasma. Before use, devices were rendered hydrophilic *via* prolonged exposure to oxygen plasma^78^.

#### μFFE device operation

Liquid-electrode microchip free-flow electrophoresis (μFFE) devices were operated as described previously^79^. Briefly, fluids were introduced to the device by PTFE tubing, 0.012"ID × 0.030"OD (Cole-Parmer) from glass syringes (Gas Tight, Hamilton) driven by syringe pumps (Cetoni neMESYS). μFFE experiments were conducted with auxiliary buffer, electrolyte, monomer reference and sample flow rates of 1000, 200, 140 and 10 μL h^-1^, respectively, for 15X reduction in buffer salt concentration for samples in PBS buffer.

Potentials were applied by a programmable benchtop power supply (Elektro-Automatik EA-PS 9500-06) via bent syringe tips inserted into the electrolyte outlets. Experiments were performed on a custom-built single-molecule confocal fluorescence spectroscopy setup equipped with a 488 nm wavelength laser beam (Cobolt 06-MLD 488 nm 200 mW diode laser, Cobolt). Photons were detected using a time-correlated single photon counting (TCSPC) module (TimeHarp 260 PICO, PicoQuant) with a time resolution of 25 ps.

#### Aggregation kinetics and sample extraction

AlexaFluor™ 488-labelled αS (100 μM) was supplemented with seed (0.5 μM) under shaking (200 rpm) at 37 °C, PBS pH 7.4 and either 1% DMSO or 50 μM molecule in 1% DMSO. Samples were extracted at the *t_1/2_* of the DMSO sample (9 hours). Fibrils were removed by centrifugation (21,130 rcf, 10 min, 25 °C) and the supernatant was then subjected to μFFE. For AlexaFluor™ 488-labelled oligomeric mixtures, auxiliary buffer comprised of 15X diluted PBS buffer, supplemented with 0.05% v/v Tween-20. Using a custom-written script, single-molecule events were recorded as discrete events using a Lee filter of 4 from the acquired photon stream as fluorescence bursts with 0.05 μs of the maximum inter-photon time and containing 30 photons minimum. Using these parameters, the single-molecule bursts and their intensities were reported as a function of device position, which could be later converted to an apparent electrophoretic mobility. Oligomer bursts were distinctly characterised by a higher photon intensity detected per molecule and a higher electrophoretic mobility than monomeric protein.

### Mass spectrometry

10 µM of preformed αS was incubated with 25 µM of molecule in 20 mM sodium phosphate buffer (pH 4.8) supplemented with 1 mM EDTA overnight under quiescent conditions at room temperature. The supernatant was removed for analysis using a Waters Xevo G2-S QTOF spectrometer (Waters Corporation, MA, USA).

### Transmission electron microscopy

10 µM αS samples were prepared and aggregated as described in the kinetic assay, without the addition of ThT. Samples were collected from the microplate at the end of the reaction (150 hours) into low-binding Eppendorf tubes. They were then prepared on 300-mesh copper grid containing a continuous carbon support film (EM Resolutions Ltd.) and stained with 2% uranyl acetate (wt/vol) for 40s. The samples were imaged at 200kV on a Thermo Scientific (FEI) Talos F200X G2 S/TEM (Yusuf Hamied Department of Chemistry Electron Microscopy Facility). TEM images were acquired using a Ceta 16M CMOS camera.

## Acknowledgements

This work was supported by the UKRI (10059436, 10061100). We thank Dr. Katherine Stott, from the Biophysics Facility, Department of Biochemistry, University of Cambridge, for her assistance in using these facilities. The authors would like to give especial thanks to Dr Laila Sakhnini for help with mass spectrometry work and Dr Heather Greer for assisting with the transmission electron microscopy (TEM) and the EPSRC Underpinning Multi-User Equipment Call (EP/P030467/1) for funding the TEM. We would also like to thank ARCHER, MARCOPOLO and CIRCE high performance computing resources for the computer time. Z. F. B. would like to acknowledge the Federation of European Biochemical Societies (FEBS) for financial support (LTF). Parts of the figures were created with BioRender.com.

## SUPPLEMENTARY INFORMATION

### Docking and Machine Learning Implementation

A full description of the initial docking approaches can be found in the previous work^1^, using AutoDock Vina^2^ and FRED^3^ docking software, but is also explained in overview here. As described in the main text, the binding site encompassing residues His50−Lys58 and Thr72−Val77 on PDB 6CU7^4^ was selected due to its propensity to form a pocket according to Fpocket^5^ software and its simultaneous mid to low solubility according to CamSol^6^ (**Figure S3**). Additionally, a key histidine residue in this site was predicted to protonate below the pH value where αS more readily aggregates (pH 5.8). A binding box was selected that had size 12 Å by 12 Å by 9 Å centred at 10.00 Å, 9.89 Å, 11.52 Å on the 6CU7 PDB, encompassing the site of interest. The target protein was left rigid, while the ligand was flexible, able to translate and rotate (including rotation of internal bonds). We prepared (added hydrogens) the target protein using Autodock tools. To increase the accuracy of the docking energy estimate, the exhaustiveness was increased compared to the default value of 8, to 20. 5 poses were output, and the best pose binding energy was selected as the binding energy label for that ligand. The choice of rigid target was made in order to decrease the computational cost of the high throughput screen of the 2 million compounds in phase 1.

Inspired by the increasing usage of consensus scoring, i.e combining multiple docking energy estimates by different docking programs, we performed docking of the 100,000 best binding molecules from AutoDock Vina, using FRED in phase 2. For each of the top 100,000 best AutoDock Vina ligands, we combine the ligand with the target into a single .pdb file, and from that supply the information of the ligand to Openye’s Spruce module to prepare an .oedu file that contains the grid position of the binding site. Then, the compound is bound to the target site and a single best pose and binding energy is output, that constitutes the FRED binding energy label for this compound. The top 10,000 are then clustered to obtain representative centroids for testing. The pipeline is modular, and it is possible to incorporate any type of docking software the user might choose. In this study we have used AutoDock Vina, which is a publicly available software that is efficient at scale, and FRED. AutoDock Vina is relatively computationally efficient at scale, and we chose to use FRED since the top scoring pose prediction of FRED has been shown^7^ to be able predict within 2Å of the native pose in 70% of examples tested. However, alternative open-source or free for academic use docking software such as rDock, LeDock and others can be used instead of FRED, with relatively little difference in performance as shown previously^8^. The performance of AutoDock Vina is comparable with other open-source software.

The code for testing the ML models on aggregation or docking data are available at https://github.com/rohorne07/Iterate. We initially tested the machine learning strategy on docking data (best R^2^ ∼0.6-0.7) before moving to experimental aggregation data (best R^2^ ∼0.2-0.3) to get an impression of the feasibility of the project, given the larger datasets available for the docking scores (**Figure S5** and **Figure S6**). The docking scores were calculated for the ‘evaluation set’, the in silico library that was used for iterative experimental screening in the main text. Both AutoDock Vina and FRED simulations were carried out on the evaluation set, giving binding scores for each molecule against the αS 6CU7 fibril structure pocket. The compound encoder was implemented as in Hie et al.^9^ to obtain representations of all the molecules. The next sections briefly summarise the functioning and output of the prediction module.

#### Prediction module

The prediction module consisted of a shallow model designed to be appropriate for small datasets and easily applicable on standard hardware available for most laboratory workers over a short timescale. As a first line test Gaussian process regression (GPR) was employed alone, following Hie et al.^9^ with training and testing carried out with cross validation on 4000 molecules from within the evaluation set. The metric used to evaluate performance in this case was the R^2^ score or coefficient of determination. This score measures the goodness of fit between a set of predictions and the ground truth values. This score ranges from 1, in a perfect fit, to arbitrarily negative values as a fit becomes worse, and is 0 when the predictions are equivalent to the expectation of the ground truth values of the training set^10^. This was compared with a naïve Bayes, which failed to score above 0 for any training set size on both docking and aggregation data.

The GPR kernel was initially the same as that utilised by Hie et al.^9^, i.e. a combination of a constant kernel and a radial basis function (RBF). Using these initial settings, R^2^ scores of ∼0.2 were obtained for the docking data. Hyperparameter optimisation yielded only marginal improvements in this performance. A selection of other kernels was tested, and all models were optimised via hyperparameter tuning before implementation, but most did not offer an improvement in performance. The Matérn kernel, a generalisation of the RBF with an extra parameter controlling the smoothness of the function, did however show a marginal improvement. These flexible functions are the most likely to be able to fit shallow energy minima problems such as those encountered here. The R^2^ scores were still low, especially for smaller training sets as would be available from experiment, but represented a viable starting point.

At this point a 2-layer model was applied. This reflected the strategy used by Hie et al. ^9^ in fitting a Gaussian process regressor (GPR) to the residuals of another model, in that case a multi-layer perceptron (MLP). An MLP did not show a dramatic improvement over the GPR alone both in that work or when tested with the docking scores here, however a random forest regressor (RFR) with stacked GPR did show a further improvement both in terms of the R^2^ (∼0.6-0.7) and the quality of the molecule sets predicted during the simulation, as can be seen in **Figure S6**.

This set up gave improved results in both R^2^ and hit rate, while retaining an easy to implement and efficient model. The average Pearson’s coefficient of correlation ranged between 0.25 and 0.3 for both the coupled (GPR+RFR) and uncoupled models (RFR alone), which while modest matched the values obtained by Hie et al. during their testing. RFR was more demanding computationally, but given the small size of the experimental training sets in this scenario this was not a hindrance.

A simulation was created to mimic how the experimental cycle of testing might work using the docking scores as a surrogate for aggregation data. In the simulation, a random subset of 100 molecules was selected and the model trained on these molecules and their binding scores. The resultant model was then used to predict binding scores for the remaining molecules and rank them using a combination of the predicted value and the associated uncertainty value. The top 100 were then selected and their binding scores added to the training set as would occur in the experimental scenario, and this process was repeated 10 times. The ideal scenario would be that molecule sets with improved mean binding energy relative to the mean of the test set would be selected, and that selections would improve as the training set expanded, and this is what is shown in **Figure S5** (though improvement is not drastic as further data is added, possibly due to the relative ease with which strong dockers are selected).

Different uncertainty penalties were tested during this process. We found that a low uncertainty penalty produced better results by removing the most overconfident predictions without placing too many limitations on the model. At the early stages most predictions with low uncertainty were those with predicted binding scores close to the mean of the training set. An excessive uncertainty penalty during these stages would cause the model to only predict molecules that it was confident in, which were also likely to be mild.

The same process was utilised using different parts of the molecular feature set (the latent vector consists of a tree vector representation of clusters within a molecule, plus a graph representation of the molecule), and it was found that GPR performance metrics were better when using the molecular graph alone compared with using the entire representation. In general, it is to be expected that fitting fewer features to a predicted value is easier for a regressor to achieve and so higher scores are obtained. However, a better average R^2^ score across the data set does not necessarily lead to a better result in terms of the actual molecules picked, and we found using the full representation led to more hits being identified (**Figure S5**).

A snapshot of the results of this testing is shown in **Figures S5** and **S6**>. **Figure S5** demonstrates 2 points: the performance was slightly improved using the Matérn kernel in place of the RBF kernel both in terms of overall hit selection and performance improvement with increasing training set size, and the full-length molecular representation gave a significant boost in terms of number of hits selected vs the truncated representation, despite lower R^2^ scores. These results also provided some evidence that Gaussian process learning might work reasonably effectively even in this data sparse scenario albeit at a modest level. It was expected that fitting experimental data would prove more challenging, however, and so a boost in performance was sought for that would not compromise the simplicity of the model, through use of the coupled RFR-GPR model. Correlation values of 0.6-0.7 were obtained using this set up on docking energies and a large portion of the dataset (4000 molecules), and this fell to between 0.2 and 0.3 for the aggregation data (**Figure S1**), which while low was encouraging given the much smaller dataset and noisier data.

---

**Table S1:** Parameters used in QSAR model optimisation. (**A**) Models such as LR and MLP were trialled with their default parameters either alone or in conjunction with a GP, but showed poor performance so were not further investigated. (**B**) GP and RF models were the best performing and so were subjected to hyperparameter optimisation via grid search cross validation using the R^2^ score as the optimisation metric. The best performing parameters are shown. The performance of these models is shown in **Figure S1**.

| Model | Parameter |  |
| --- | --- | --- |
|  | *Parameter label* | *Parameter value* |
| LR | fit_intercept | True |
| LR | copy_X | True |
| LR | n_jobs | 1 |
| LR | positive | False |
| MLP | hidden_layer_sizes | 100 |
| MLP | activation | relu |
| MLP | solver | adam |
| MLP | alpha | 0.0001 |
| MLP | batch_size | auto |
| MLP | learning_rate | constant |
| MLP | learning_rate_init | 0.001 |
| MLP | max_iter | 200 |
| MLP | tol | 0.0001 |
| MLP | warm_start | False |
| MLP | early_stopping | False |
| MLP | beta_1 | 0.9 |
| MLP | beta_2 | 0.999 |
| MLP | epsilon | 1×10<sup>-8</sup> |
| MLP | n_iter_no_change | 10 |

LR = linear regressor, MLP = multi layer perceptron

B
| Model | Parameters |  |  |
| --- | --- | --- | --- |
|  | *Kernel (if applicable)* | *Parameter label* | *Parameter value* |
| GP | Constant | length_scale | 1.0 |
| GP | Constant | length_scale_bounds | fixed |
| GP | RBF | length_scale | 1.0 |
| GP | RBF | length_scale_bounds | fixed |
| GP | Matern | length_scale | 1.0 |
| GP | Matern | length_scale_bounds | fixed |
| GP | Matern | nu | 1.5 |
| RF | n/a | n_estimators | 950 |
| RF | n/a | criterion | squared_error |
| RF | n/a | max_depth | 50 |
| RF | n/a | min_samples_split | 2 |
| RF | n/a | min_samples_leaf | 2 |
| RF | n/a | min_weight_fraction_leaf | 0.0 |
| RF | n/a | max_features | log2 |
| RF | n/a | max_leaf_nodes | None |
| RF | n/a | min_impurity_decrease | 0.0 |
| RF | n/a | bootstrap | False |
| RF | n/a | oob_score | False |
| RF | n/a | warm_start | False |
| RF | n/a | max_samples | None |

GP = Gaussian Process, RF = random forest, RBF = Radial Basis Function

**Table S2:** Clinical and neuropathological characteristics of synucleinopathy and non-synucleinopathy brain tissue samples used in the study.

| Brain Sample | Source | Gender (M/F) | Age at Death (years) | Disease duration (years) | Postmortem interval (h) | Primary diagnosis | Additional diagnosis |
| --- | --- | --- | --- | --- | --- | --- | --- |
| DLB1 | Ghetti | M | 81 | N.A. | 20 | Diffuse Lewy body disease | Senile changes, Cerebrovascular disease |
| DLB2 | BSFRC | M | 70 | 6 | N.A. | Lewy body Dementia | Senile changes, Cerebrovascular disease |
| DLB3 | BSFRC | M | 75 | 4 | N.A. | Lewy body Dementia | Senile changes, Cerebrovascular disease |
| MSA1 | BSFRC | F | 62 | N.A. | N.A. | Multiple system atrophy | N.A. |
| MSA2 | Ghetti | F | 71 | N.A. | N.A. | Multiple system atrophy | Senile changes, Cerebrovascular disease |
| MSA3 | Ghetti | M | 52 | N.A. | 3 | Multiple system atrophy | Senile changes, Cerebrovascular disease |
| CBD | Ghetti | F | 51 | 10 | 9.6 | Corticobasal Degeneration | N.A. |

**Figure S1.**
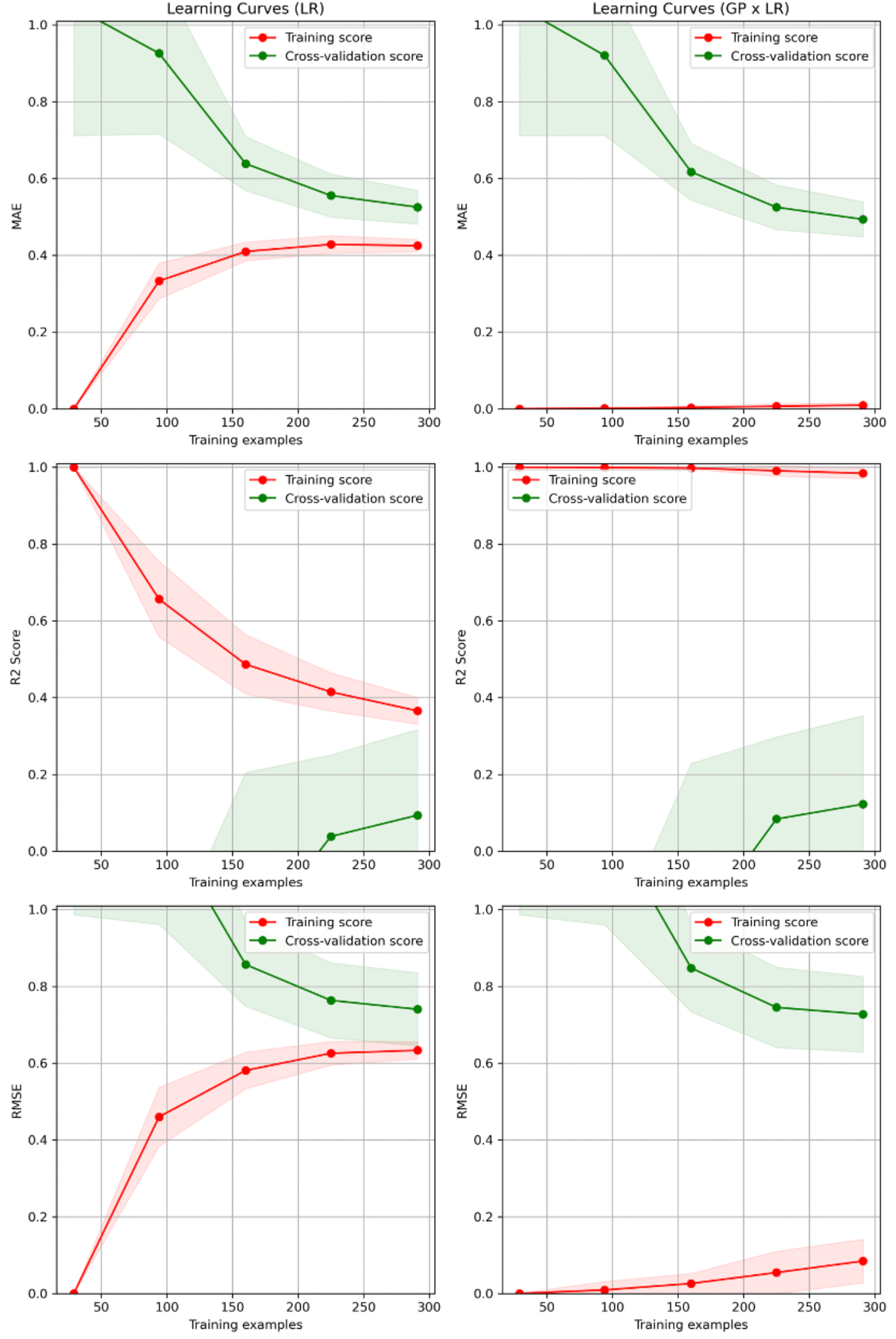
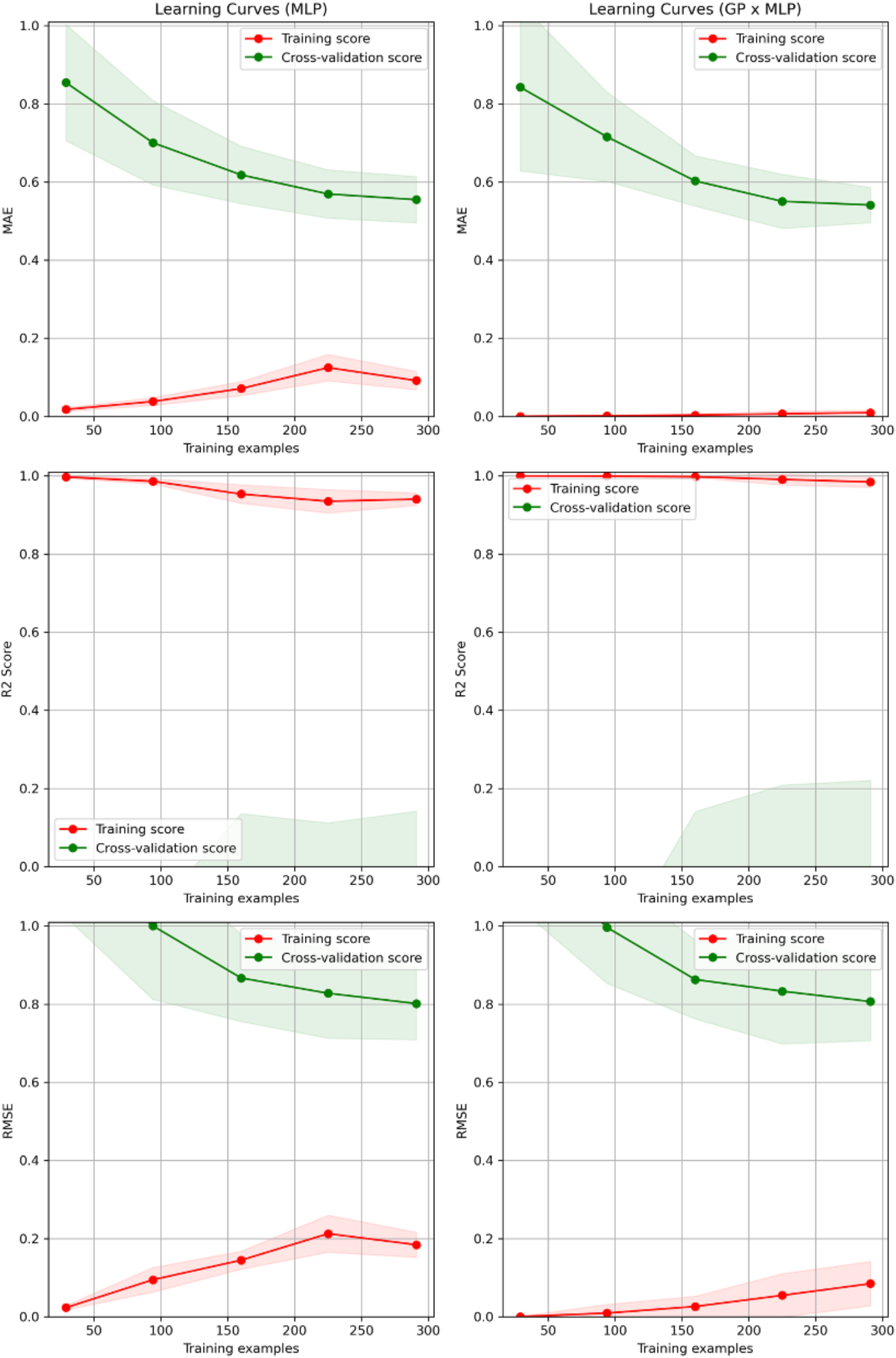
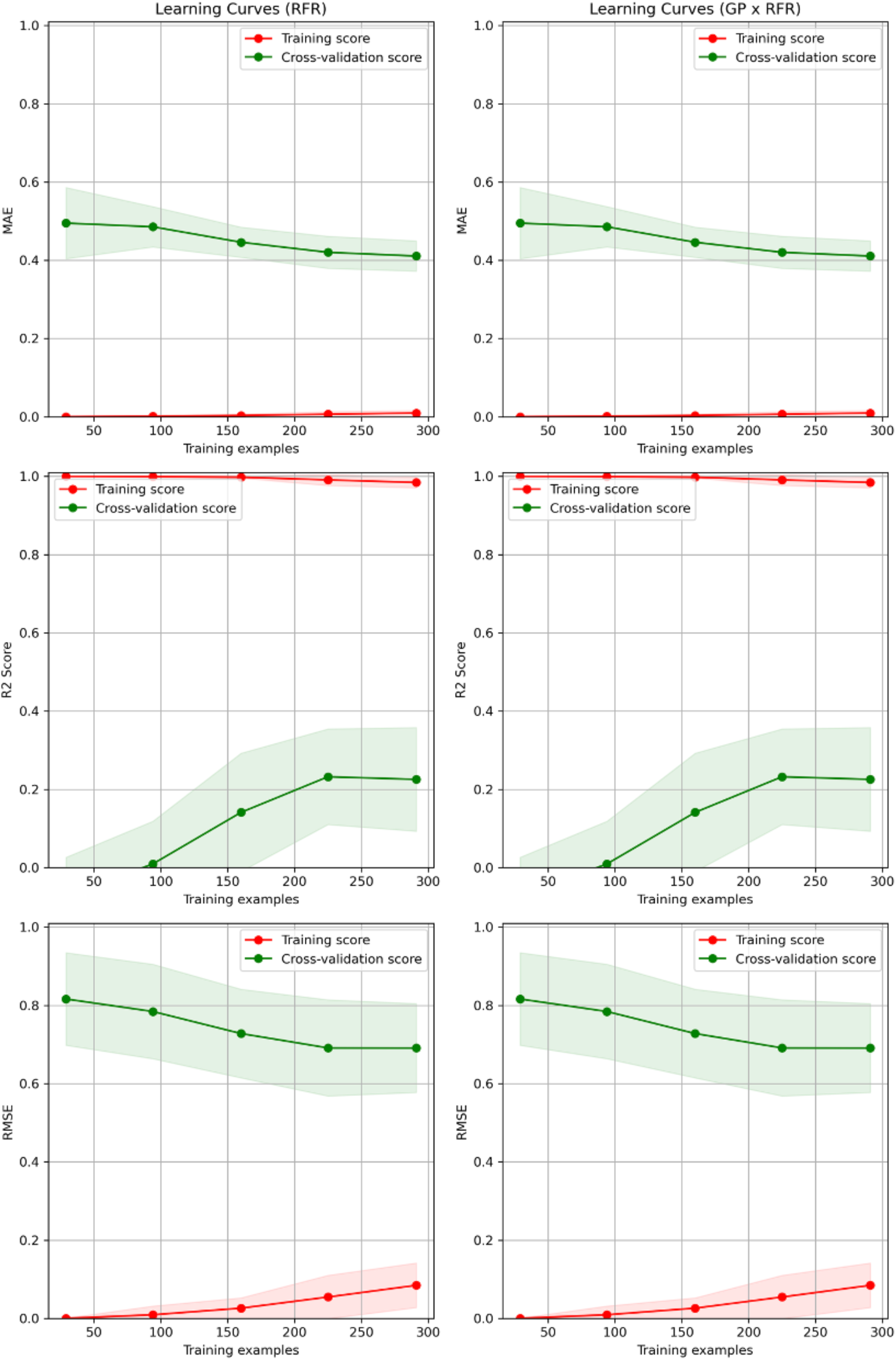
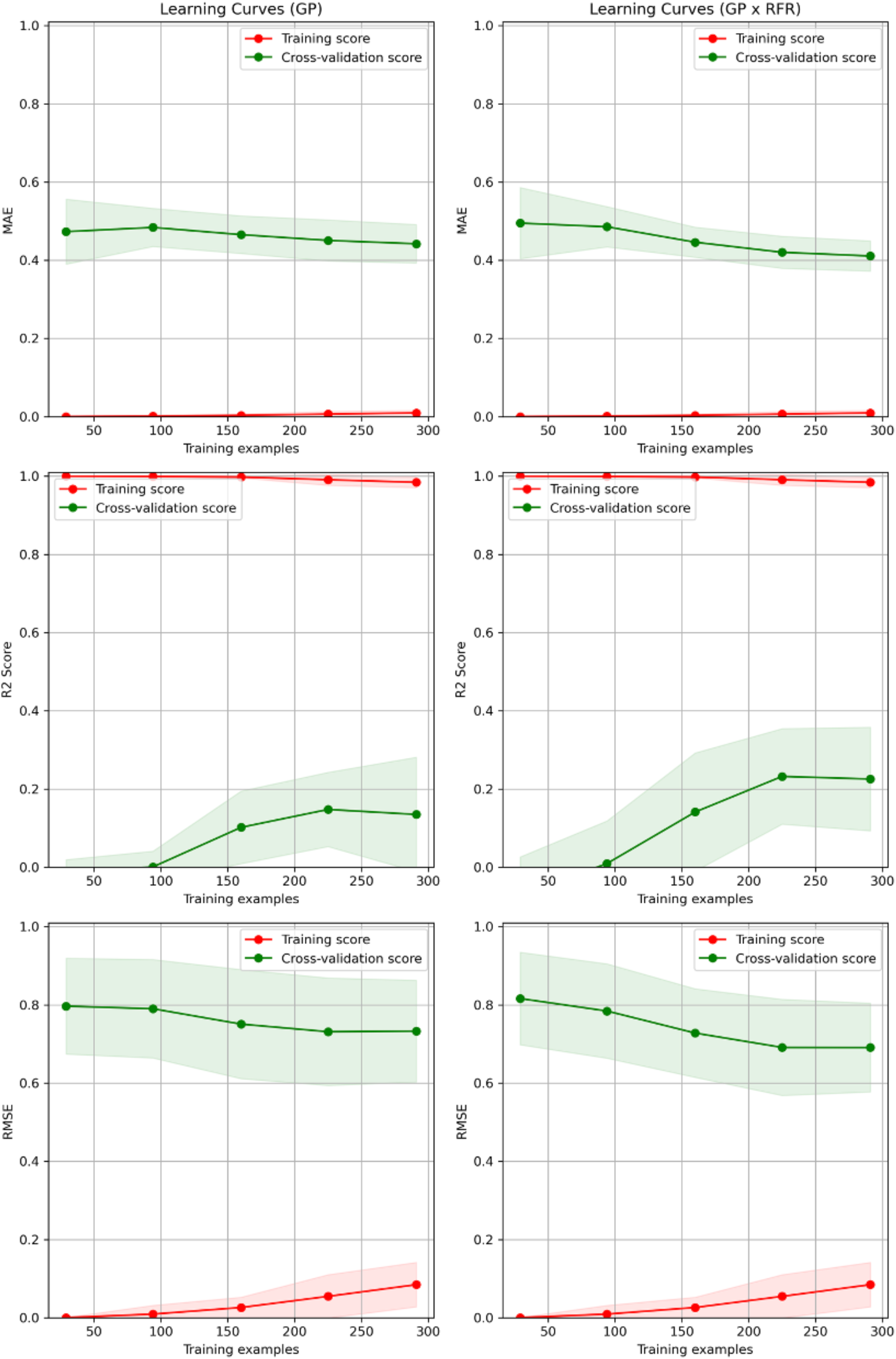
MAE, RMSE and R^2^ for different models trained on the latent features of the variational autoencoder and the aggregation data. The y-axis reports the respective scoring metric, and the x-axis the number of molecules included in the training set, of a total sample of 360 molecules. In each case, the performance of the model in isolation is shown in the left column, while the performance of the model when used in tandem with the GPR fitted to the residuals of the first model is shown in the right column. The labels are as follows: LR = linear regressor, GP = Gaussian process, MLP = multilayer perceptron, RFR = random forest regressor. Model parameters were chosen using a grid search of possible parameters while cross validating on 5 stratified K folds of the aggregation data, and selecting the parameters that gave the best performance in terms of R^2^ score. The parameters for the models shown here are displayed in **Table S1.**

**Figure S2.**
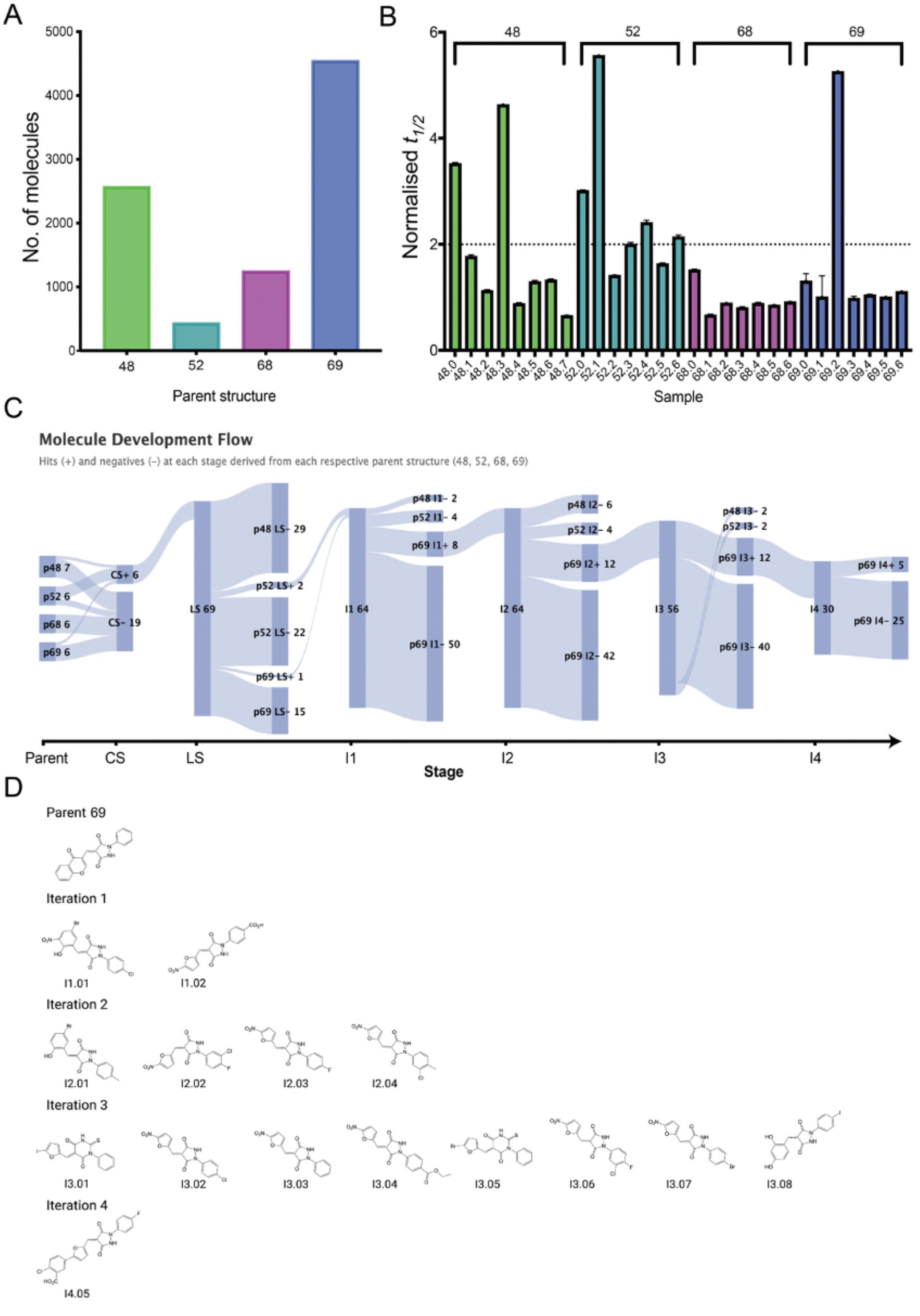
Summary of the molecules described in this work. **(A)** Number of molecules derived from 1 of the 4 docking hits (48, 52, 68, 69) within the evaluation set (see Figure 1). There were more structures derived from molecules 69 and 48 compared with molecules 68 and 52. **(B)** Normalized half time of aggregation (*t_1/2_*) for the 25 molecules in the close similarity docking set (25 µM), i.e. those closely related (Tanimoto similarity > 0.5) to the 4 molecules in the docking set (labelled as 48.0, 52.0, 68.0 and 69.0 on the x-axis). Hits were defined as molecules that more than double *t_1/2_*, as indicated by the horizontal line that marks 2 times the half time (y-axis) in the absence of the molecules. Some derivatives of molecules 48, 52 and 69 showed good potency, in particular 48.3, 52.1 and 69.2, but these effects were outstripped by future hits such as I4.05 which yielded the same effects at 50 fold lower concentration. **(C)** Flow chart of molecule hits (+) and negatives (-) in the project starting from the close search (CS), moving to the loose search (LS) then iterations 1, 2, 3 and 4 (I1, I2, I3, I4). Each branch is labelled with the molecule source (e.g. parent 48 = p48) whether it was a hit or a negative, and the number of molecules in the branch. Attrition reached its highest point at the loose search before gradually improving with each subsequent iteration. Iteration 4 is included but not directly comparable as a model was trained on the lower dose inhibition for this step. **(D)** Structures of the most potent hits at each stage, which flatlined aggregation at 25 µM, all of which were derived from p69. The structures gradually converged as the core pyrazolidine-3,5-dione structure and RHS aromatic ring were largely retained (with some exceptions for ring expanded derivatives in iteration 3) with addition of electron withdrawing groups to the benzene ring. The LHS was altered more significantly, replacing the parent bicyclic system with substituted furans, which were further elaborated in iteration 4 with an additional benzoic acid group.

**Figure S3.**
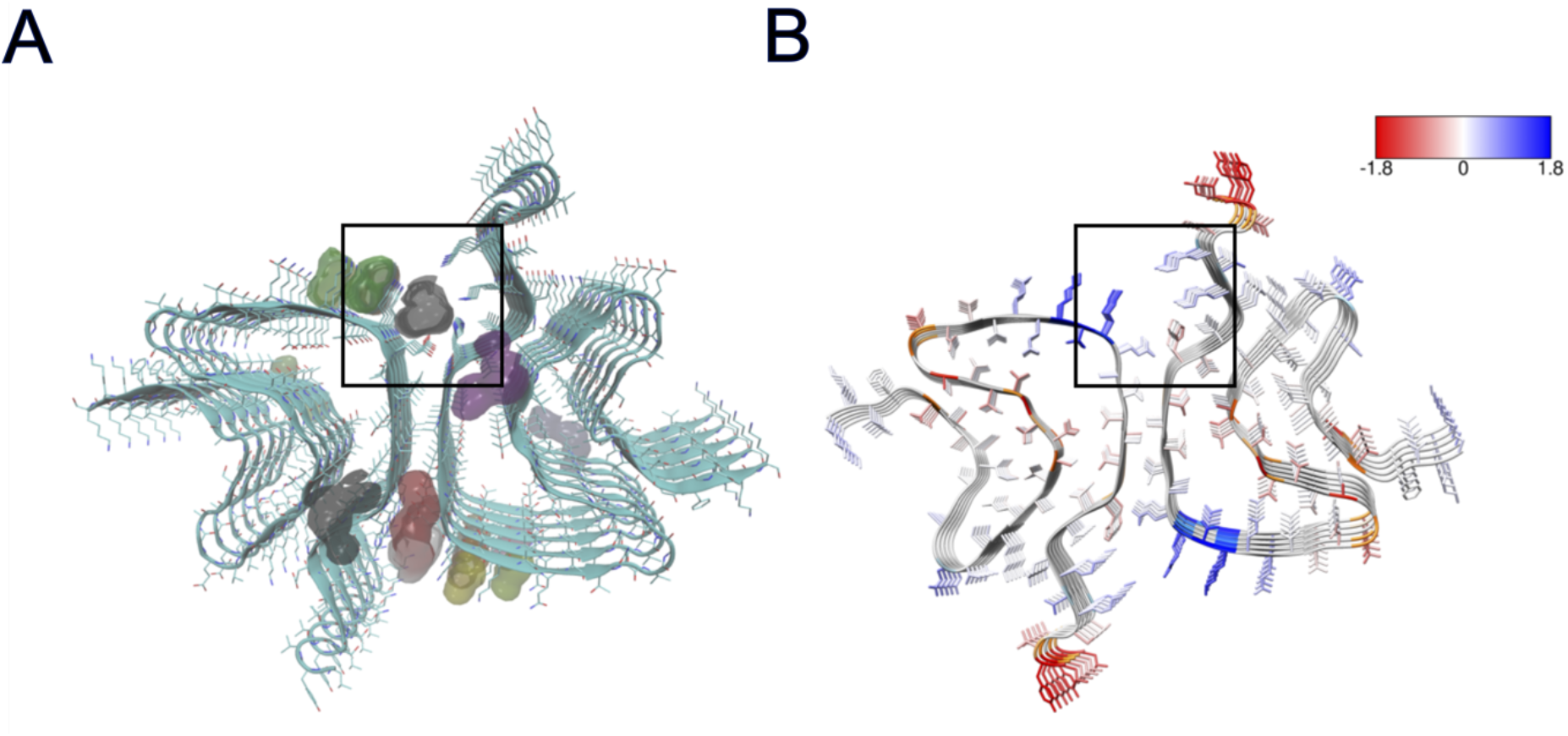
Volume and solubility based binding site prediction on polymorph 6CU7^4^. **(A)** Cavity based binding site prediction based on Fpocket^5^. **(B)** Solubility based binding site prediction based on CamSol^6^. The black box outlines the region encompassing key residues His50 and Glu57 where both cavity propensity is high and solubility is medium-low.

**Figure S4.**
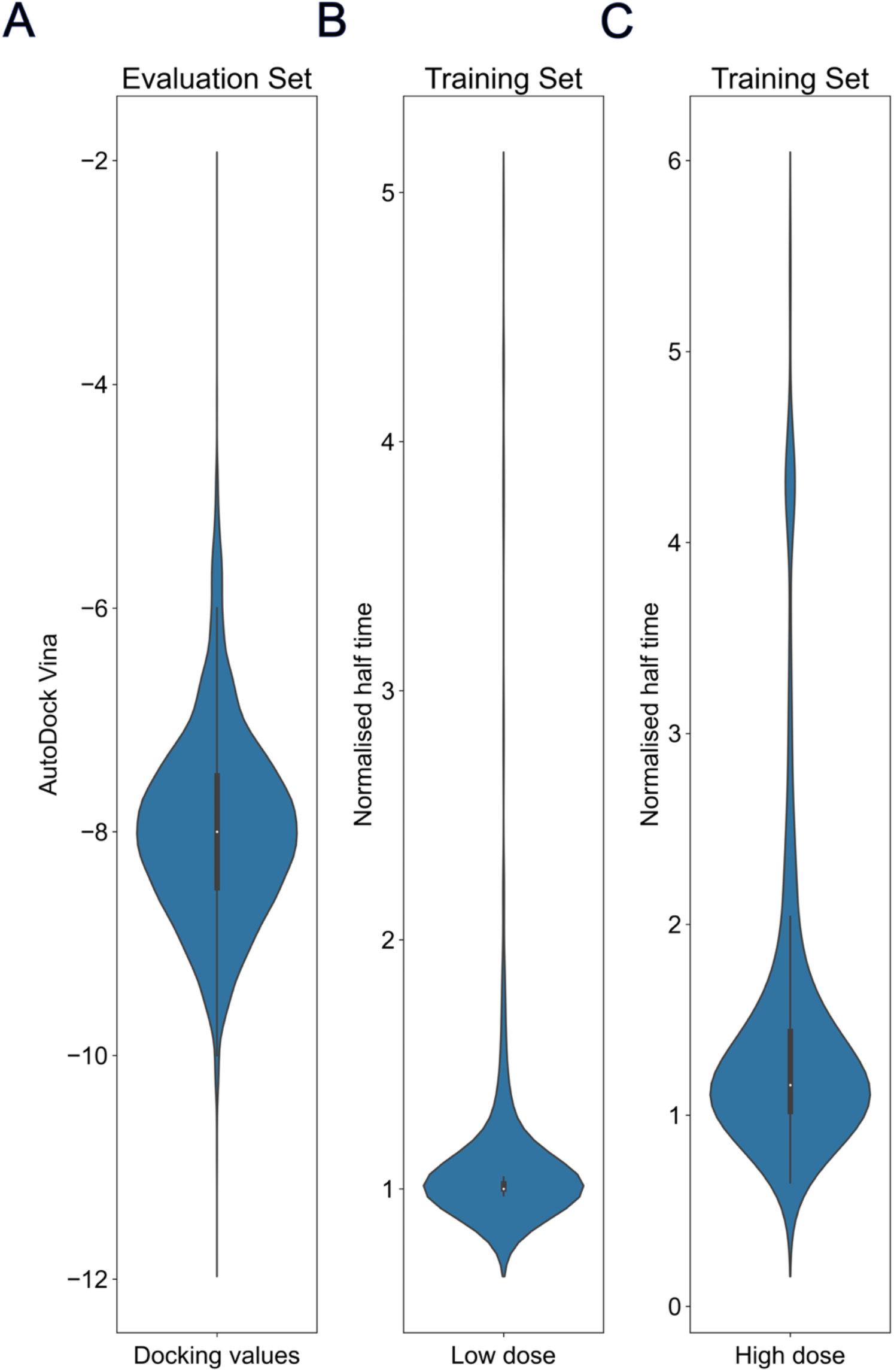
Distributions of the datasets used. **(A)** AutoDock Vina binding energies (kcal mol^-1^) for the evaluation set (∼9000 molecules). The values are narrowly distributed between −6 and −10 kcal mol^-1^ as the dataset consists of 4 key structures predicted to have good binding. Normalised half times of aggregation at **(B)** 3.12 µM and **(C)** 25 µM for the whole training set (∼400 molecules), including docking molecules and initial similarity searches and after all iterations had been added. The high dose was used for training in iterations 1-3 and the low dose for iteration 4.

**Figure S5.**
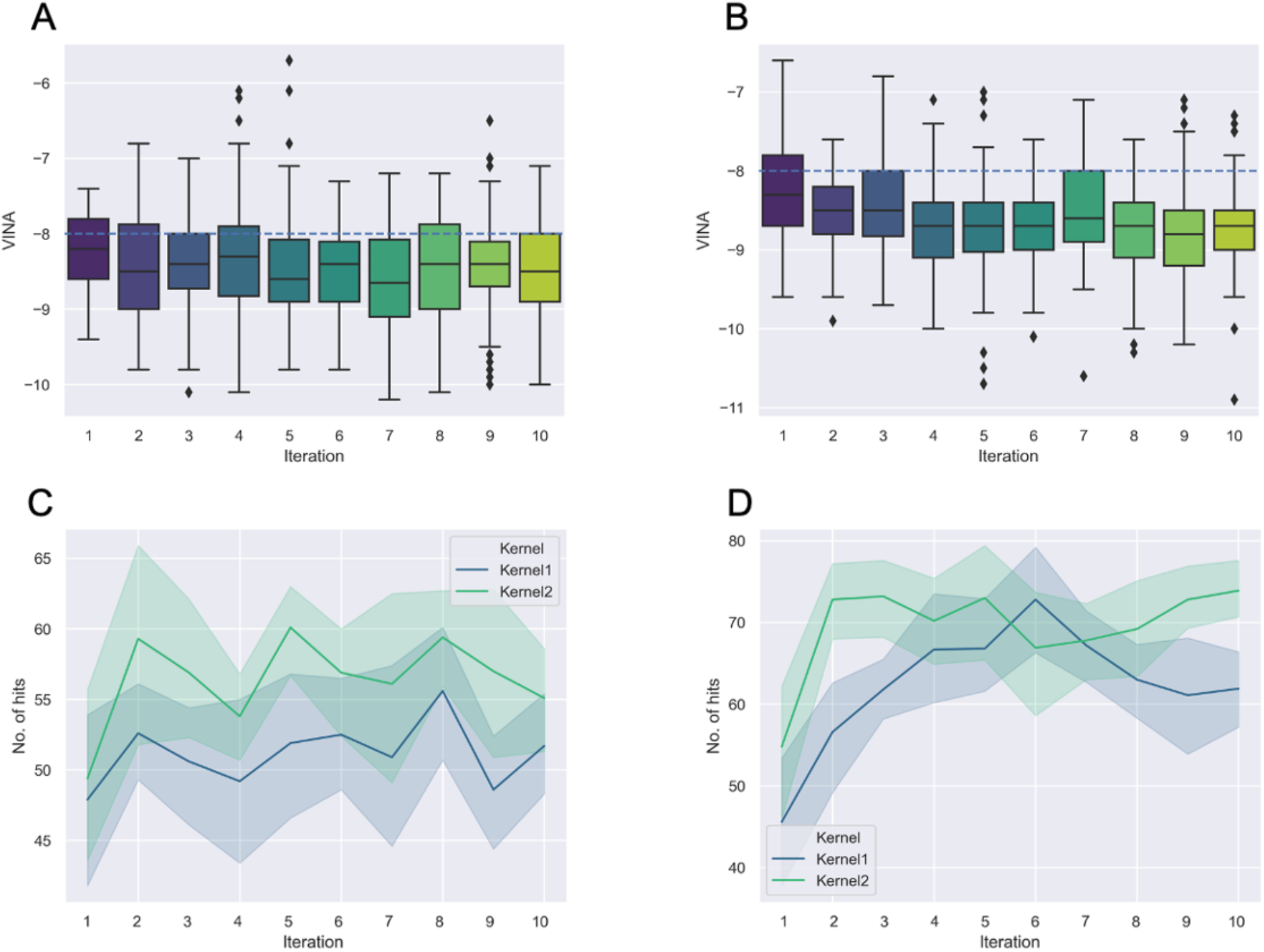
A simulation of the experimental scenario using docking energies as a proxy for aggregation experiments. **(A)** Starting from a single random sample, the GP with RBF kernel was tested. AutoDock Vina binding energies in kcal mol^-1^ are plotted against iteration number. Each boxplot visualises the distribution of binding scores for the top 100 molecules predicted by the algorithm at each iteration. The dotted line indicates the mean binding energy of the test set. **(B)** Same process as in panel A, but employing the GP with a Matérn Kernel. **(C)** Aggregated average number of hits out of the top 100 predicted molecules from 10 different random starts of the process shown in panels A and B for the RBF kernel (Kernel 1, in blue) and the Matérn kernel (Kernel 2, in green). A hit was taken as a molecule falling in the lower quartile of the test set distribution (<-9 kcal/mol). Results were obtained using the half-length representation of the molecules. **(D)** Same process as described in panel C, but employing the full-length molecule representation.

**Figure S6.**
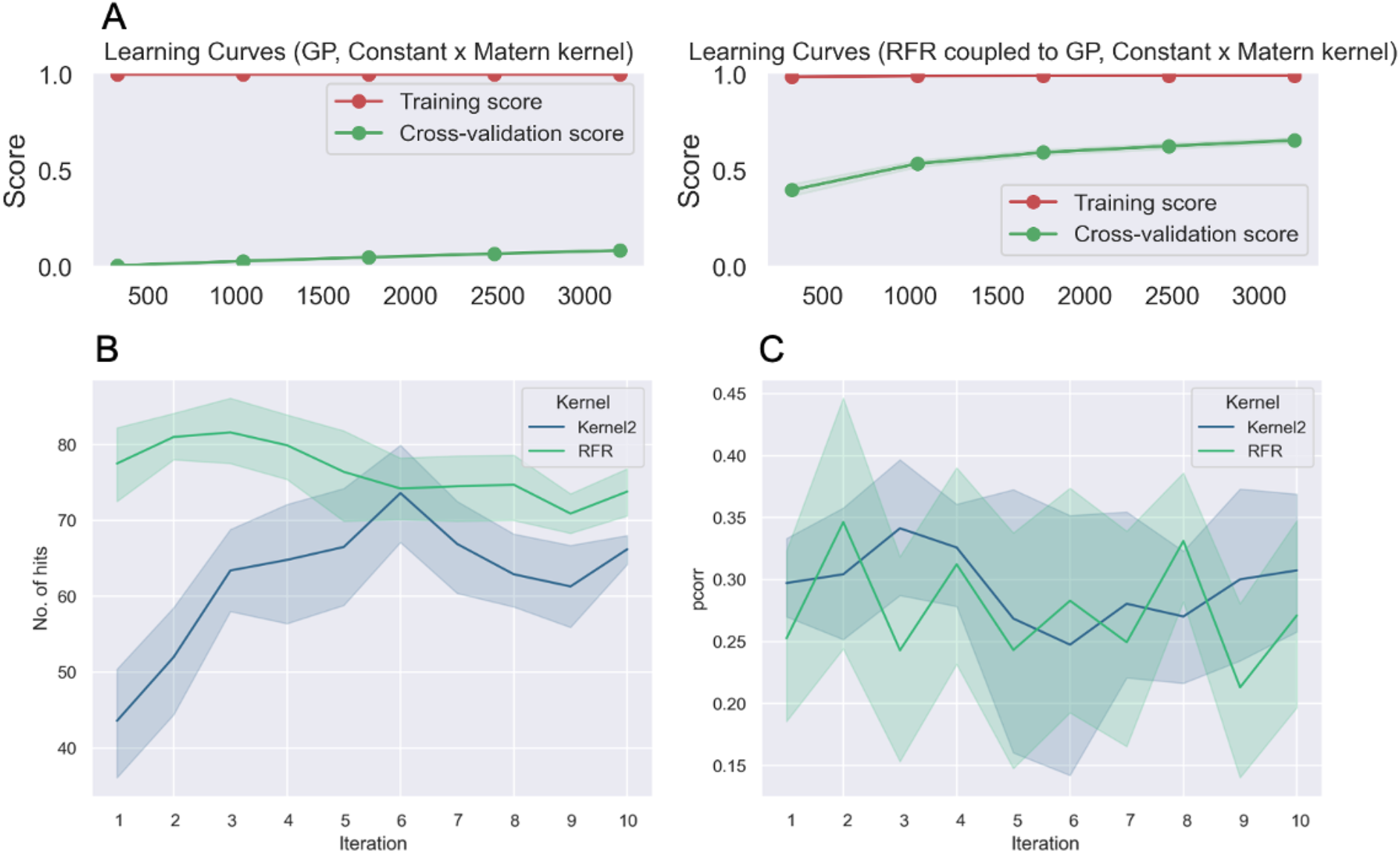
Performance of the RFR method coupled to the Matérn kernel compared to the Matérn kernel alone. **(A)** R^2^ score with increasing training set size (up to 4000) for both models, using the full-length representation. On the left is the GP with Matérn kernel alone, and on the right is the GP with Matérn kernel + RFR. Cross validation with 10 random shuffle splits and 20% of the data randomly selected as a validation set. **(B)** Aggregated average hit data from 10 different random starts of the experimental simulation for the iterative approach, starting from 100 randomly selected molecules and successively adding the actual docking data of the predicted top 100 hits to the training set with each iteration. GP with Matérn kernel alone (Kernel 2 = Matérn) vs GP with Matérn kernel + RFR. **(C)** Average Pearson’s correlation coefficient (pcorr) between the predicted binding score values and the real scores at each iteration.

**Figure S7.**
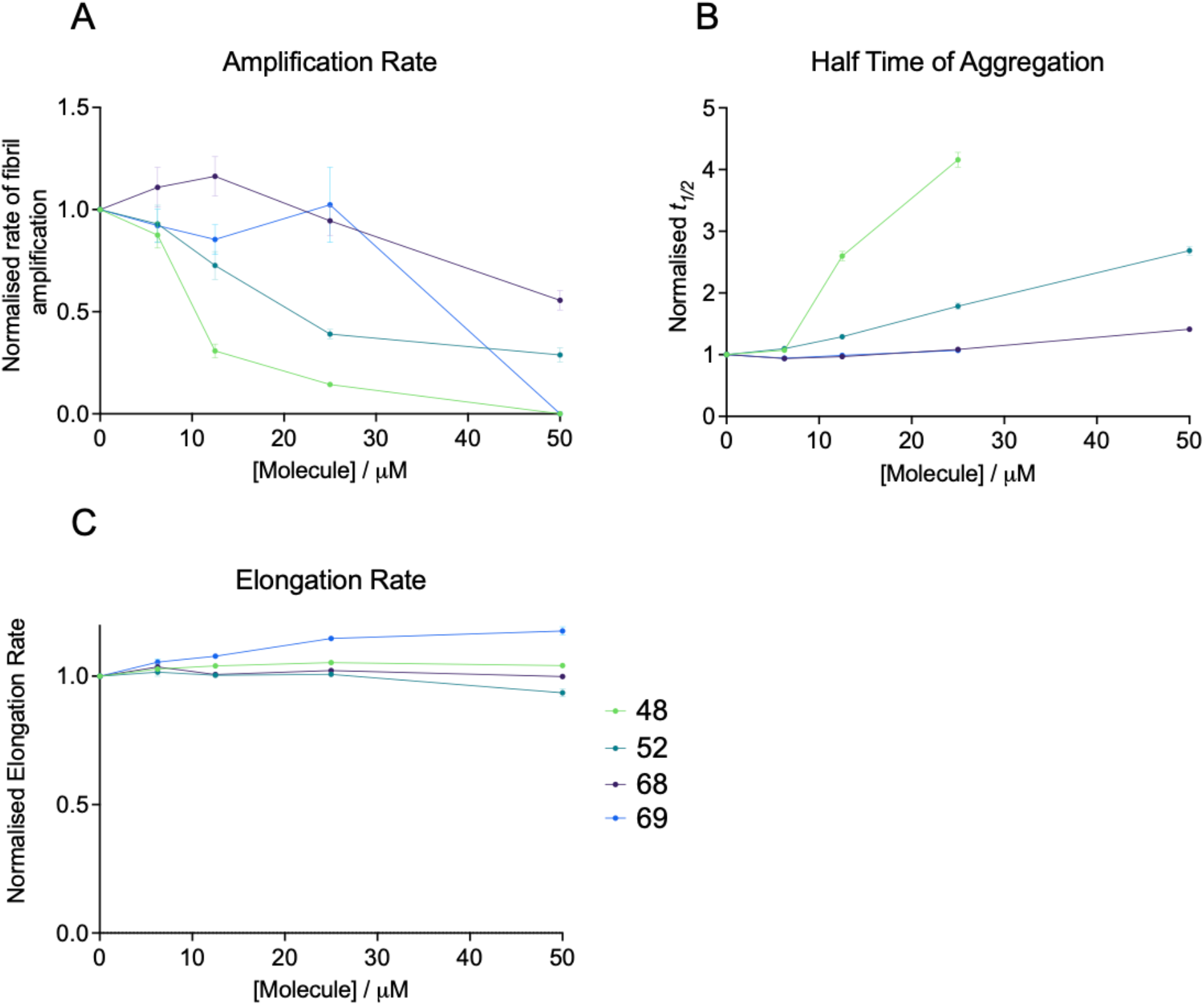
Amplification rate and half time of aggregation of αS in the presence of the 4 molecules in the docking set. **(A)** Relative rate of fibril amplification of αS in the presence of the 4 docking molecules (labeled as 48, 52, 68 and 69) in the docking set; the kinetic traces are normalised to the DMSO control. **(B)** Half times of aggregation derived from the same experiment. (**C**) Relative rate of fibril elongation normalised to the DMSO control. The amplification rate (A) and half time of aggregation (B) were tested in the machine learning method as parameters to describe the potency of a molecule. The amplification rate tends to be more affected by perturbations to the early slope of the exponential phase can have large effects on the derived rate value. The half time, although a simpler measure, is more robust and so was chosen for the machine learning approach. Data obtained from reference^1^.

**Figure S8.**
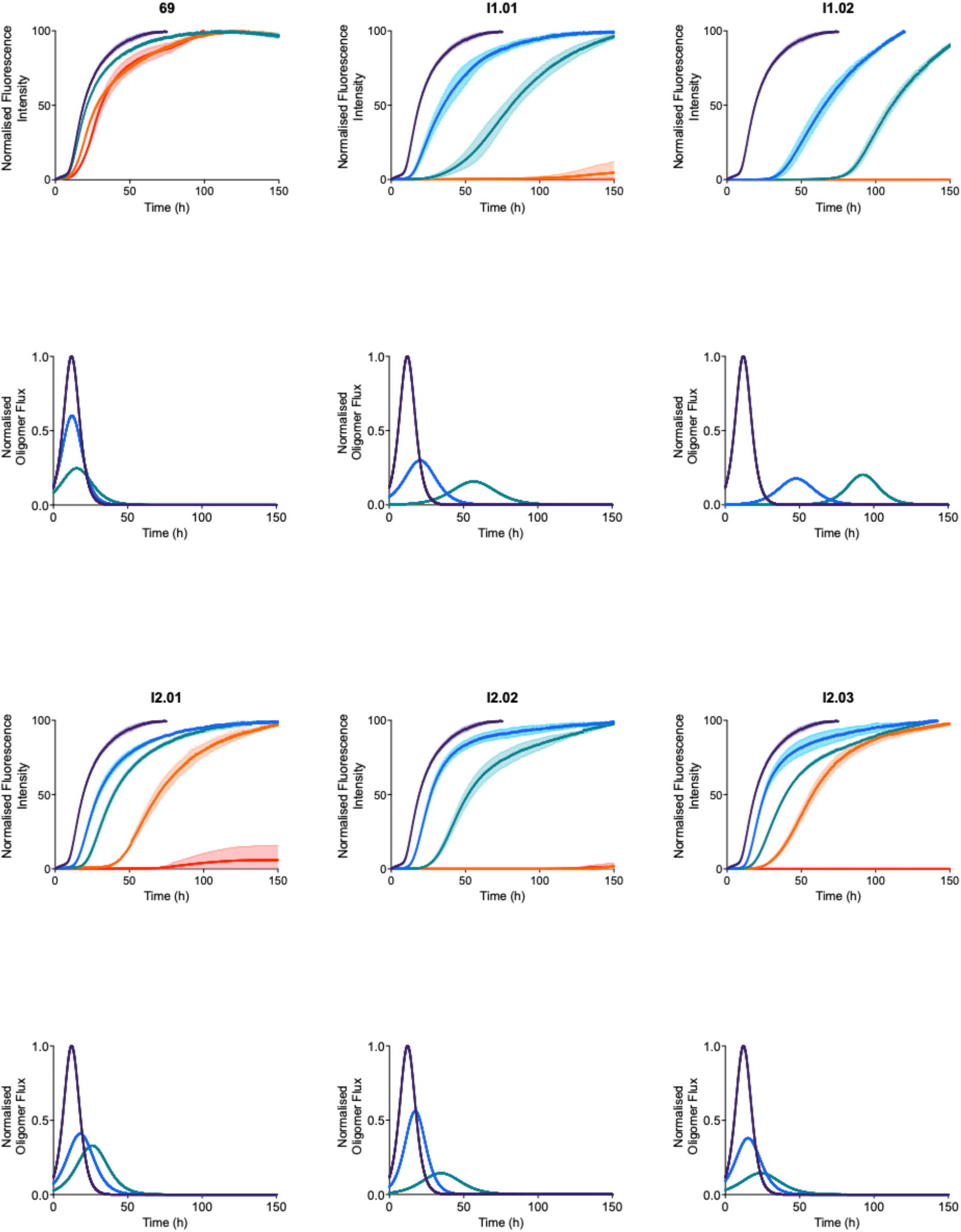
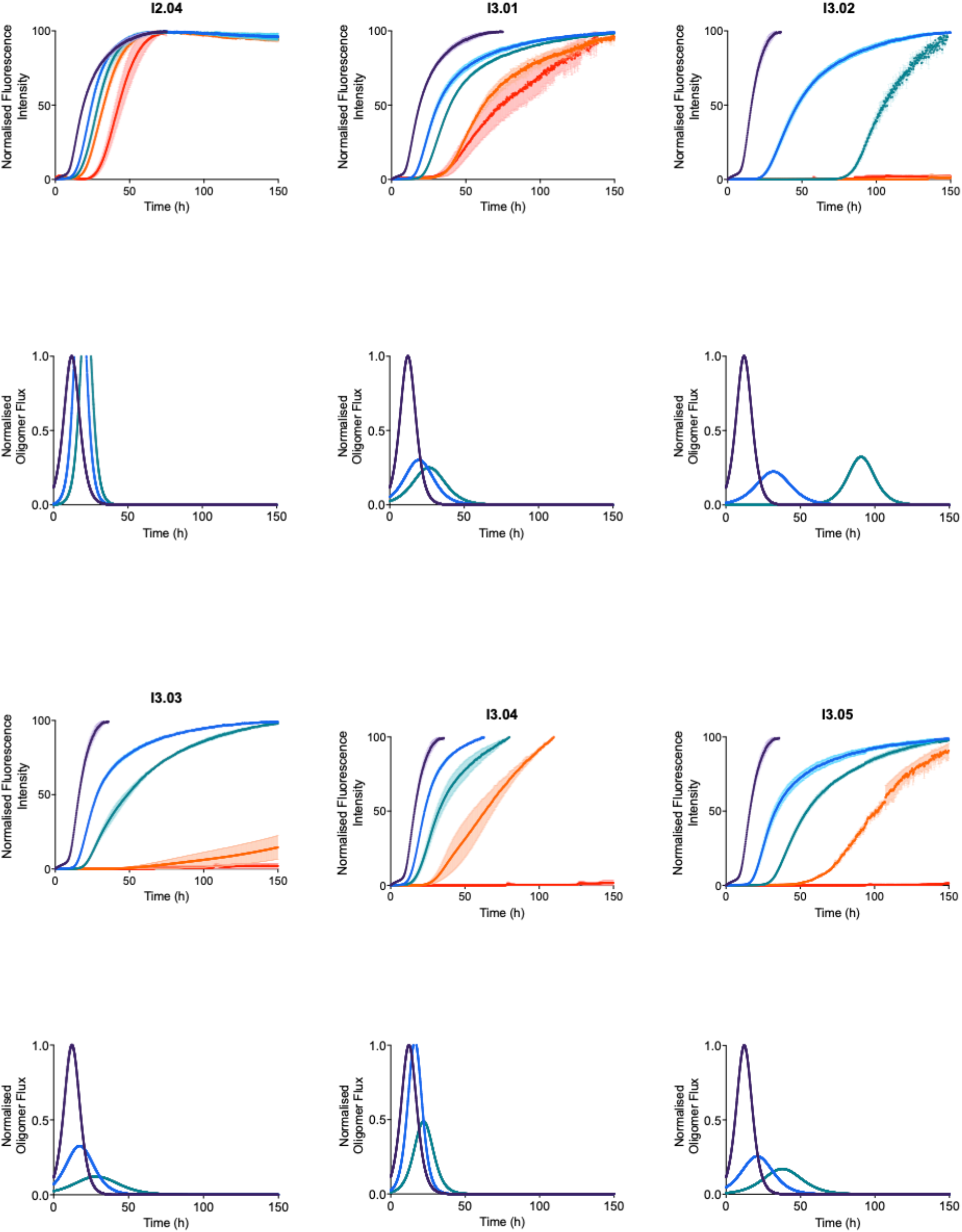
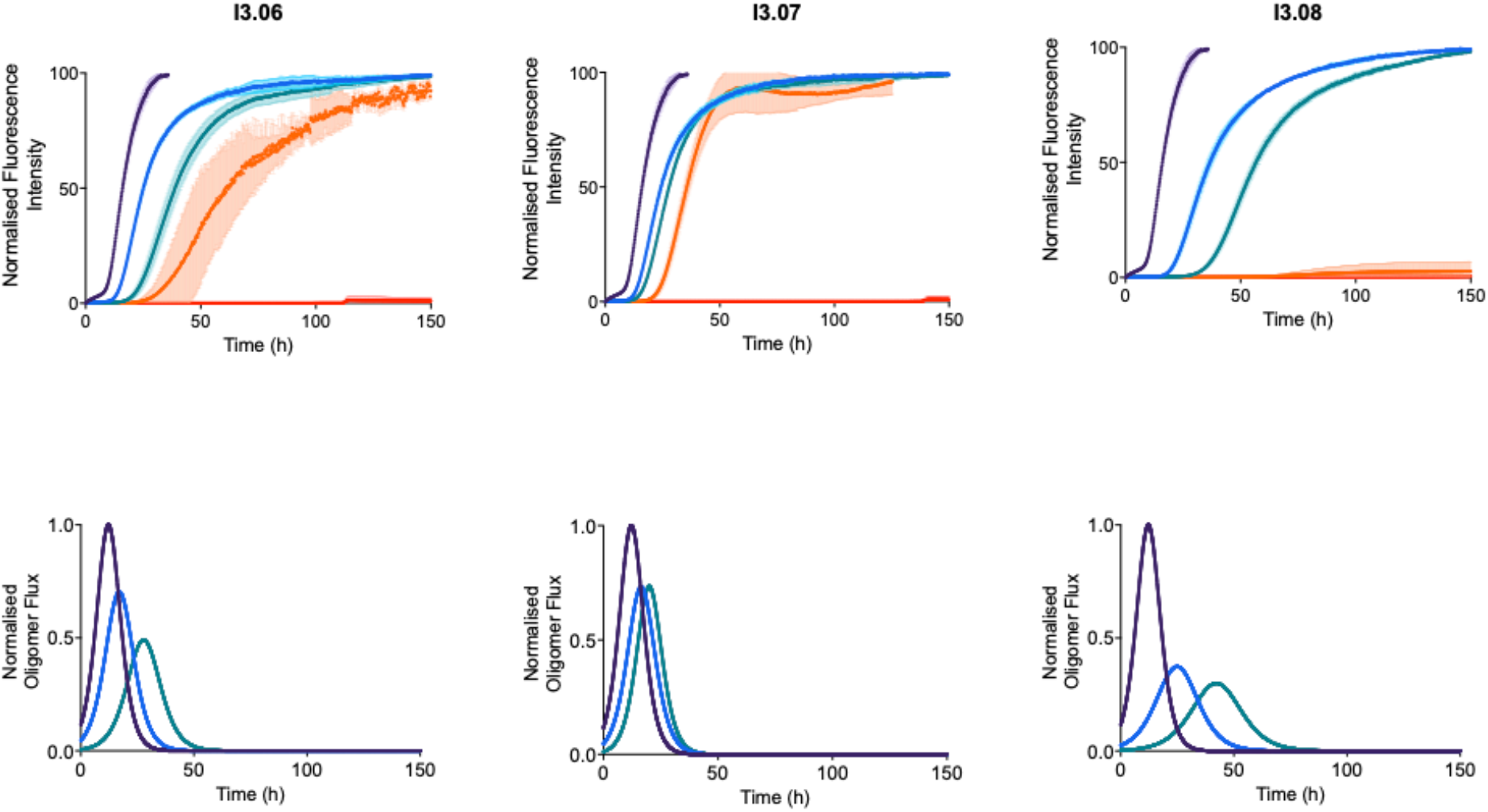
Aggregation curves (top) and oligomer flux simulations (bottom) for the most potent compounds from all of the iterations. The kinetic traces show a 10 µM solution of αS in the presence of 25 nM seeds at pH 4.8, 37 °C in the presence of molecules at 3.12 µM (blue), 6.25 µM (teal), 12.5 µM (orange) and 25 µM (red) versus 1% DMSO alone (dark purple), with endpoints normalised to the αS monomer concentration detected via the Pierce™ BCA Protein Assay at the end of the experiment. Oligomer simulations were carried out only for the lower 2 concentrations, as full aggregation curves were only consistently obtained for all molecules in the secondary nucleation assay at these concentrations.

**Figure S9.**
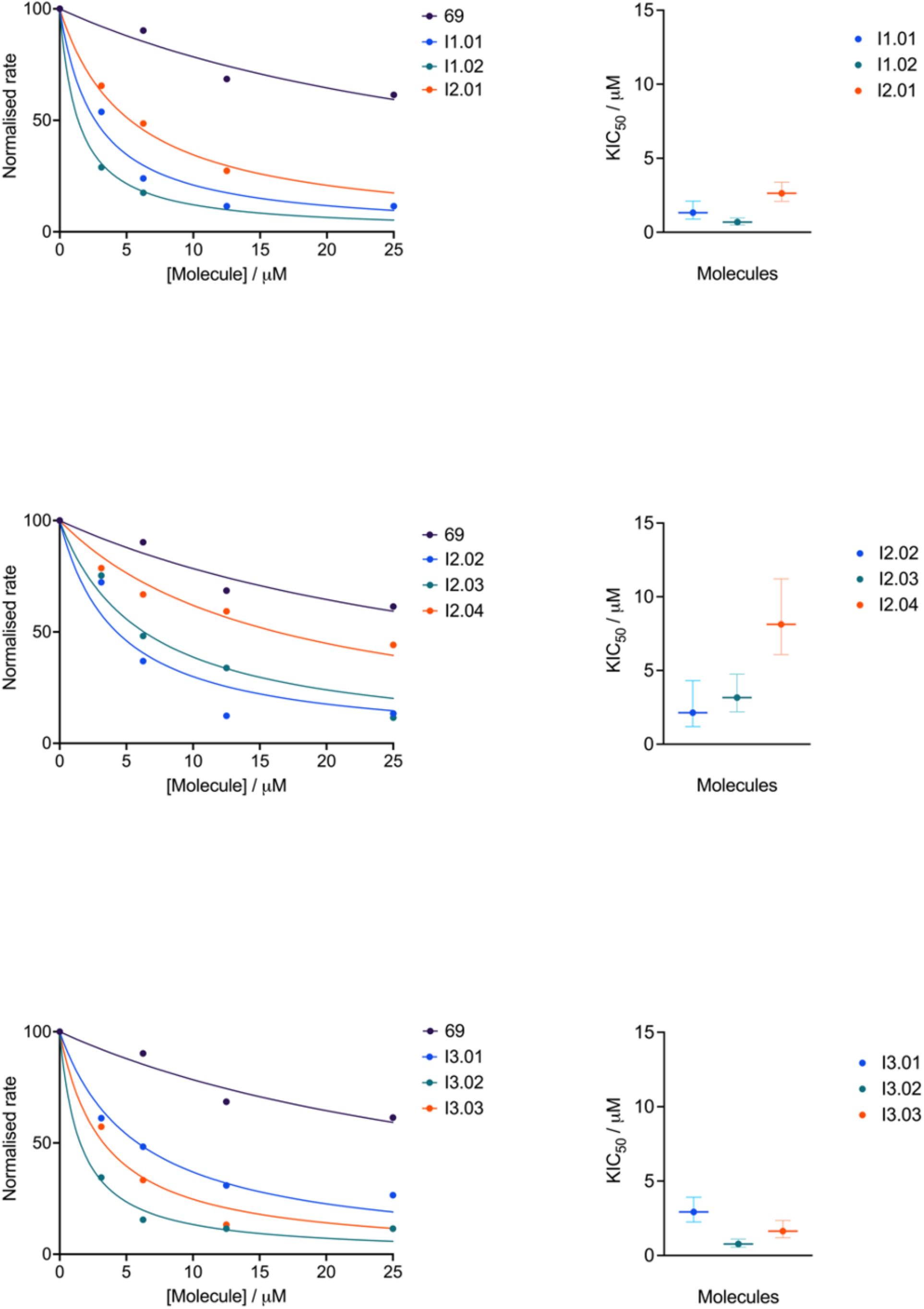
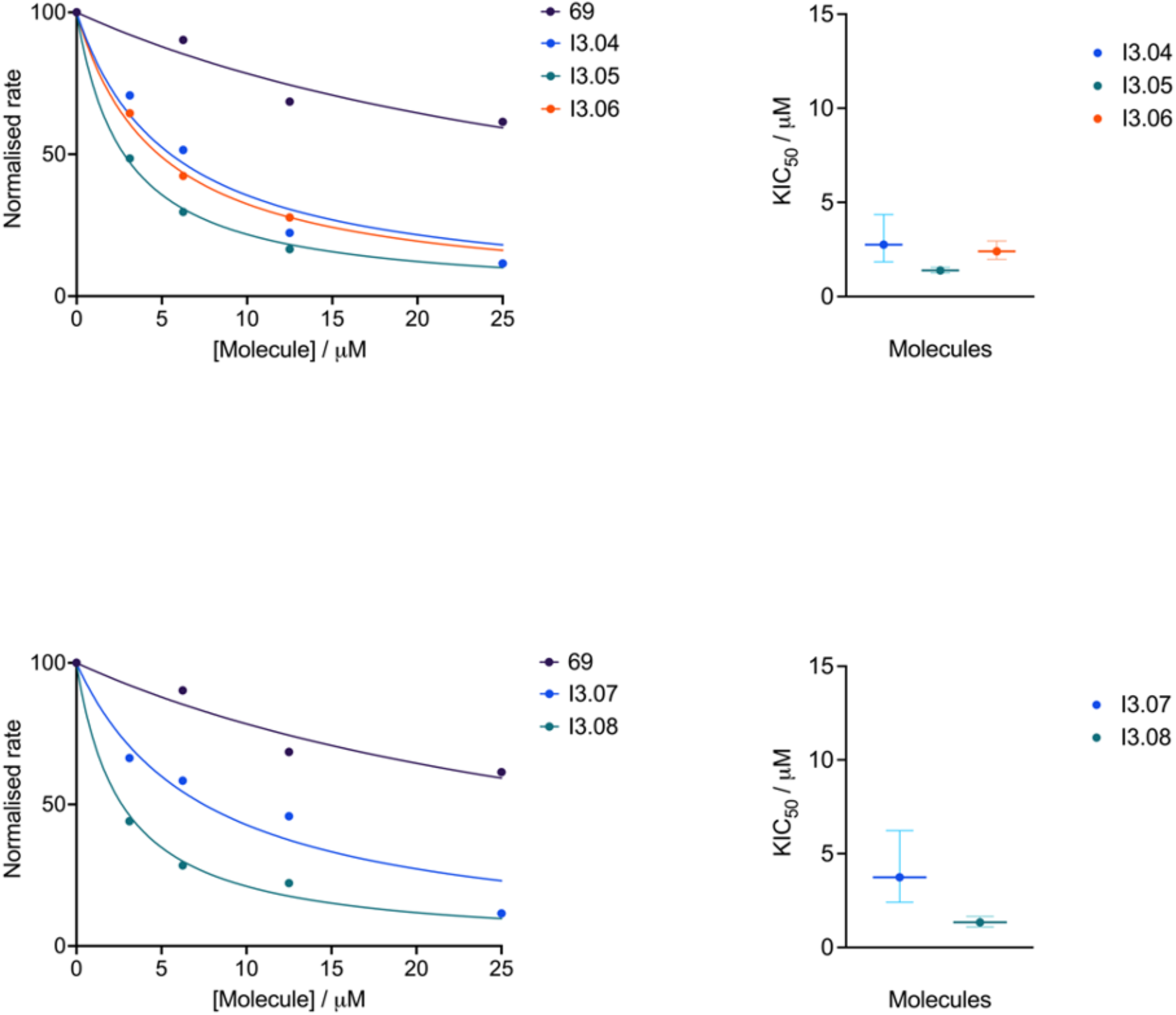
Concentration dependence of the reaction rate and corresponding 50% kinetic inhibitory concentration (KIC_50_) values for the most potent compounds. The approximate normalised rate of reaction (taken as 1/*t_1/2_*) is shown on the left for each molecule at each concentration for which a half time could be obtained. For molecules that completely inhibited the aggregation process on the timescale of the experiment, the *t_1/2_* in the presence of the highest concentration of molecule (25 µM) was taken to be the length of the experiment. The approximate rates are fitted using an [Inhibitor] vs. normalized response Hill slope. The KIC_50_ values are shown on the right with the 95% confidence interval.

**Figure S10.**
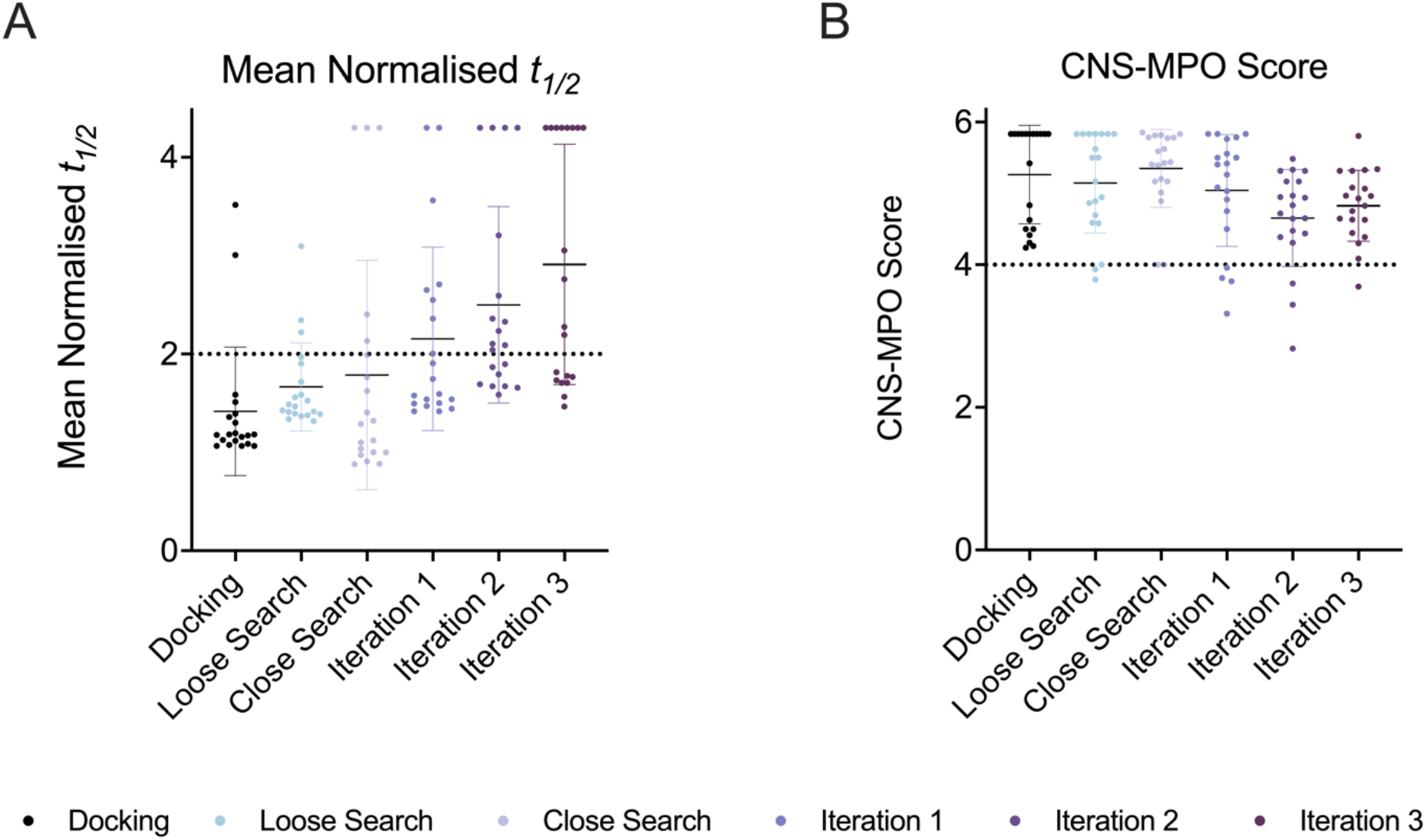
Average *t_1/2_* of aggregation and CNS-MPO scores for the top 20 molecules at each stage. **(A)** The stages are the initial docking simulation (68 molecules tested), loose search (69 molecules tested), close search (25 molecules tested), iteration 1 (64 molecules tested), iteration 2 (64 molecules tested) and iteration 3 (56 molecules tested). Molecules were tested at a concentration of 25 µM during screening. Molecules that completely prevented aggregation were assigned a *t_1/2_* value equal to the length of the experiment. **(B)** A common cut off for CNS-MPO score is 4, as indicated by the horizontal dotted line.

**Figure S11.**
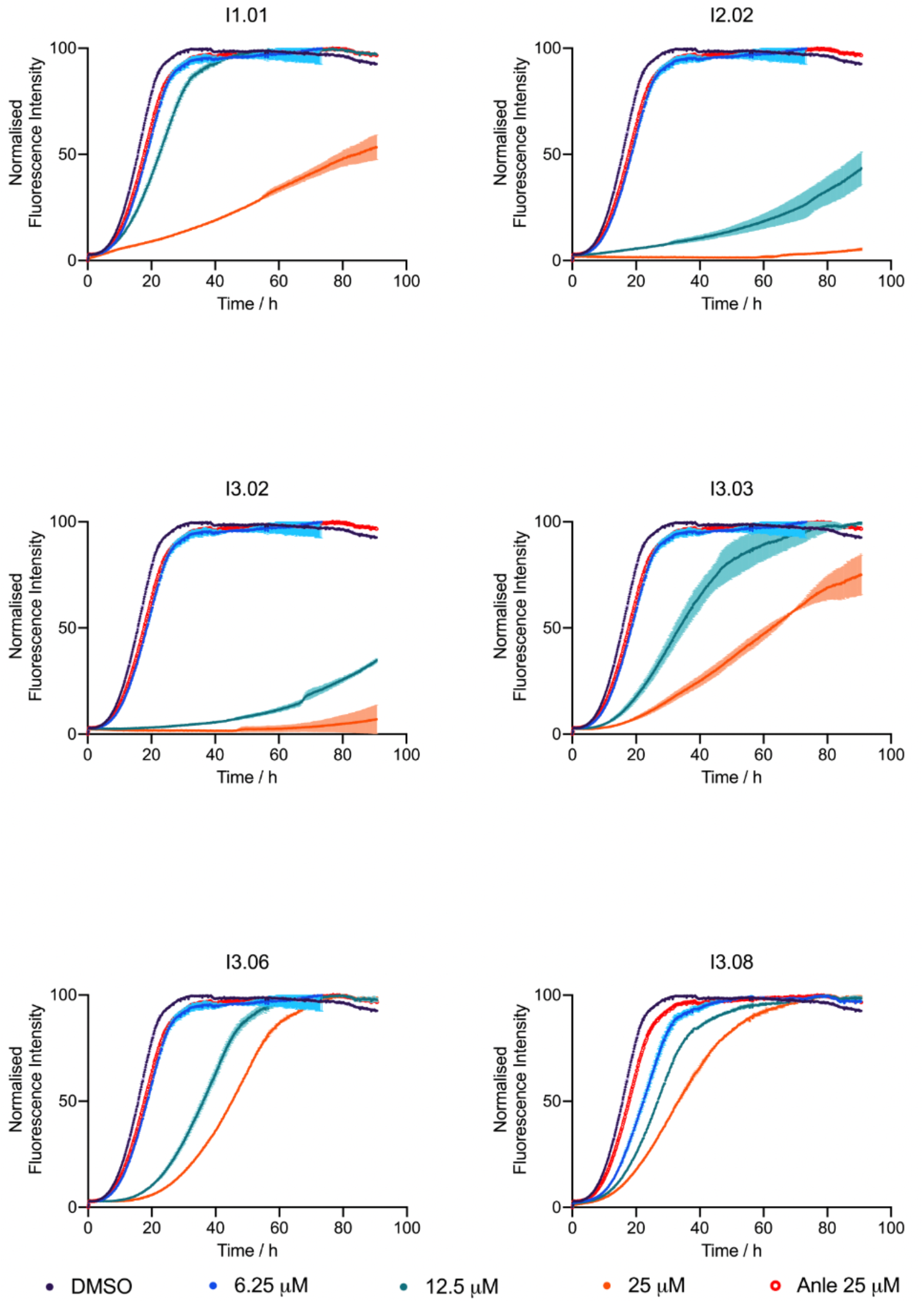
Lipid induced aggregation curves in the presence of the early hits from the project. The kinetic traces show a 20 µM solution of αS in the presence of 100 µM DMPS vesicles (monomer equivalent) at pH 6.5, 30 °C in the presence of molecules at 6.25 µM (blue), 12.5 µM (teal), 25 µM (orange) and Anle-138b at 25 µM (red circles) versus 1% DMSO alone (dark purple), with endpoints normalised to the αS monomer concentration detected via the Pierce™ BCA Protein Assay at the end of the experiment.

**Figure S12.**
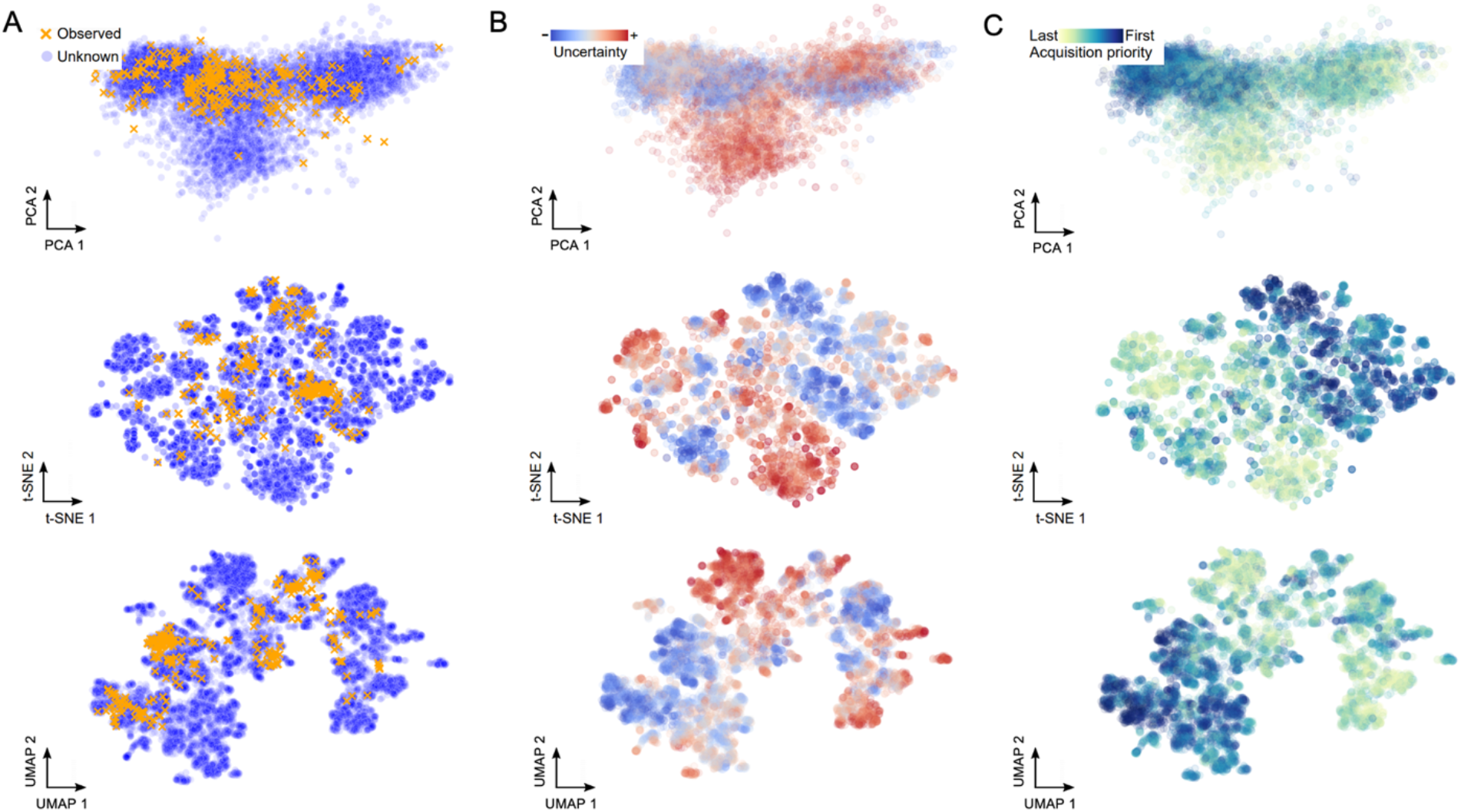
PCA, t-SNE and UMAP visualisations of the compound feature space using uncertainty. (A) From top to bottom: PCA, t-SNE and UMAP visualisations of the compound space indicating which areas of the chemical space have been explored (orange crosses) and which have not (blue circles). **(B)** GPR assigned lower uncertainty (blue) to regions of the chemical space near to the observed data and high uncertainty (red) to areas which were further away. **(C)** The lower uncertainty compounds were prioritised (dark blue) during acquirement ranking.

**Figure S13.**
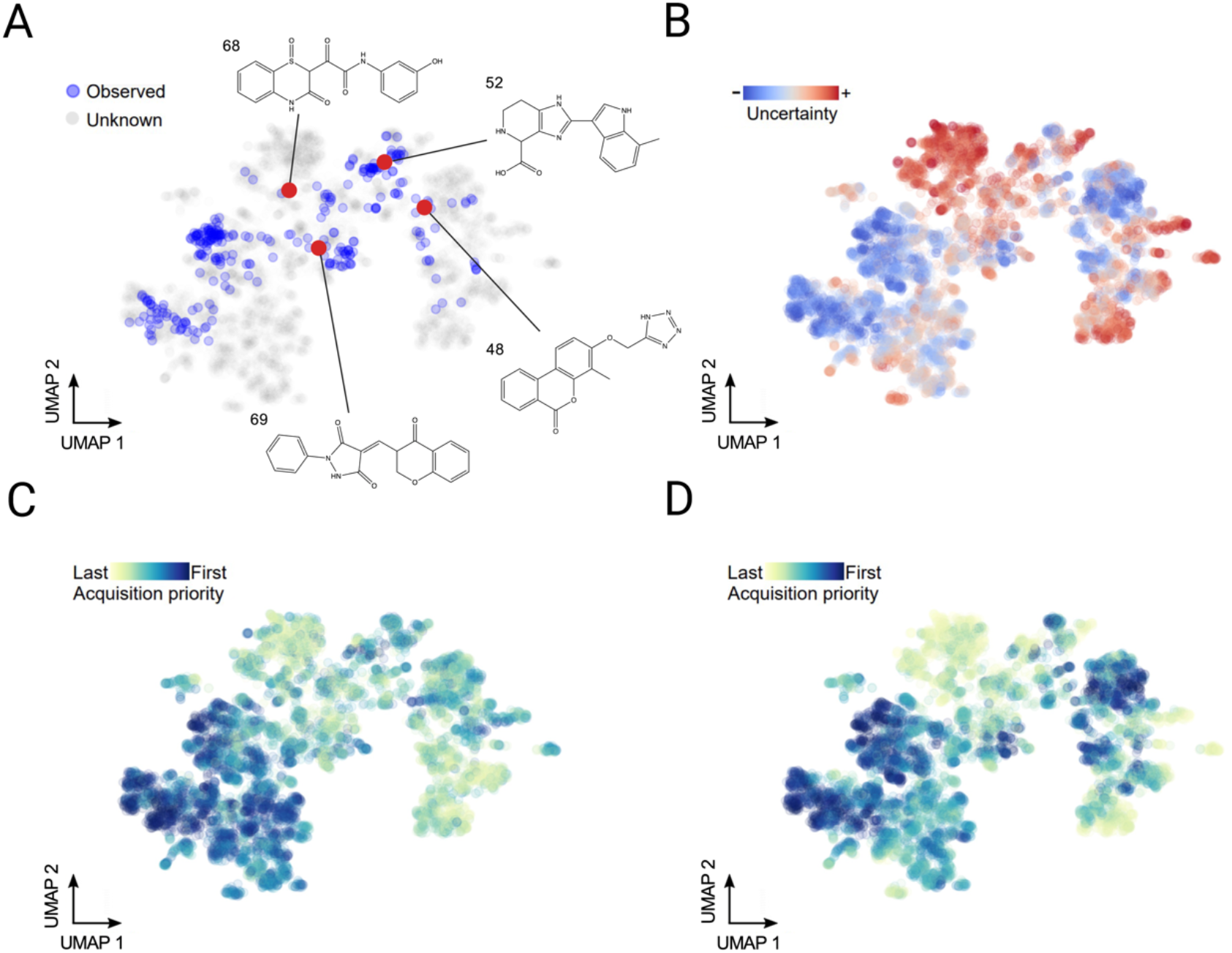
UMAP visualisation of the compound feature space using uncertainty. **(A)** The visualisation indicates the molecules in the chemical space that have been tested over the course of the project (blue circles) starting from the 4 initial docking molecules (red circles) in the docking set, and the relative positioning of the parent structures in this space. **(B)** GPR assigned lower uncertainty (blue) to regions of the chemical space near to the observed data and high uncertainty (red) to areas which were further away. **(C)** Acquirement ranking with a low uncertainty penalty. The lower uncertainty compounds were prioritised (dark blue) during acquirement ranking. **(D)** Acquirement ranking with a high uncertainty penalty.

**Figure S14.**
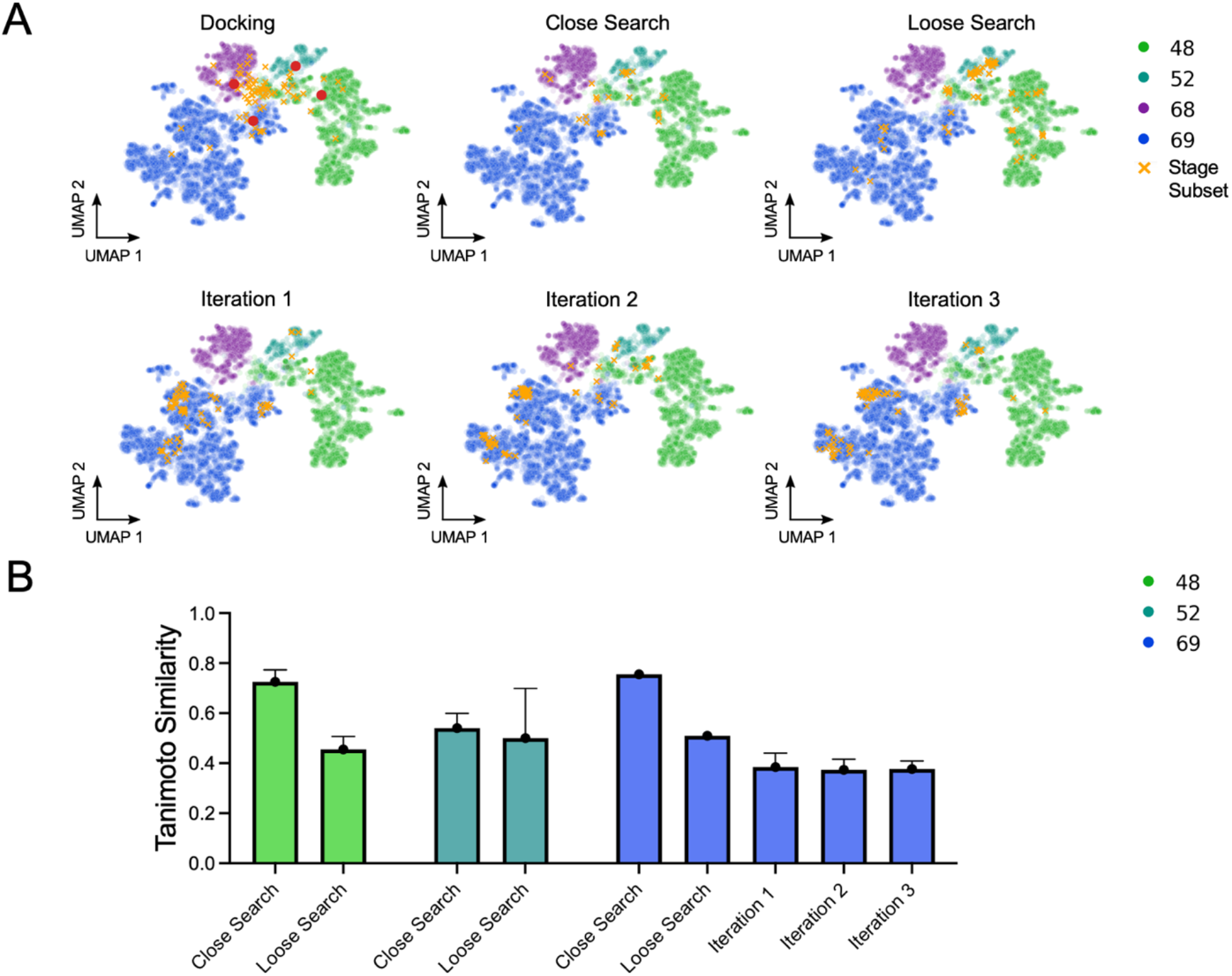
Analysis of the structural changes in the compound optimisation. **(A)** UMAP visualisation of the compound space indicating how the positioning of each new molecule subset (orange crosses) changed at each stage of the project as well as how the chemical landscape was split between the parent molecules (different colours) The locations of the parent molecules are also indicated in the ‘Docking’ pane (red circles). **(B)** Average Tanimoto similarity of the hit molecules to their respective parents at each stage of the project. At iterations 1, 2 and 3 all of the hits were derived from molecule 69, albeit with lower similarity than any of the previous stages. Molecule 68 failed to produce any hits outside of the parent molecule.

**Figure S15.**
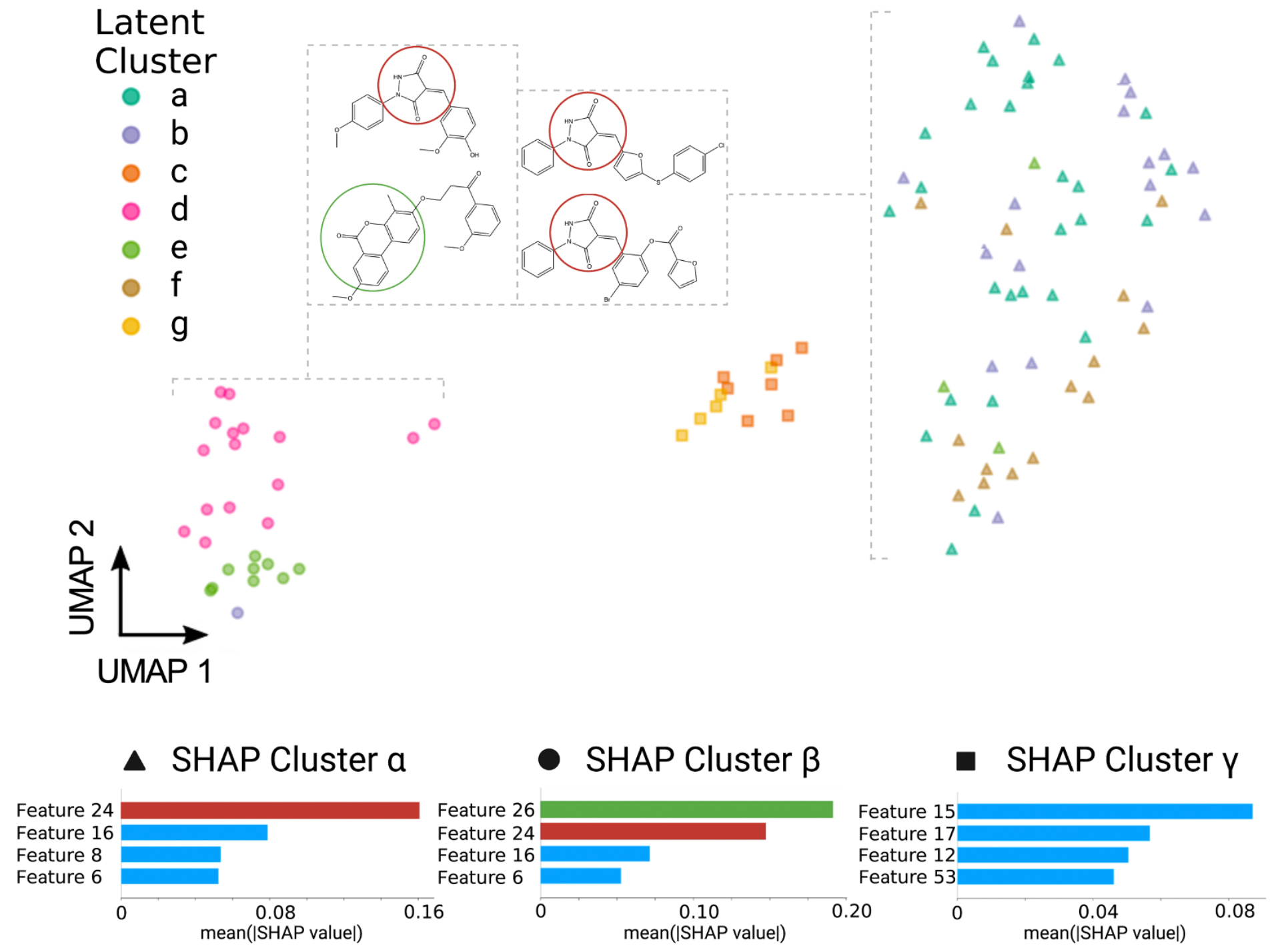
Clustering molecules based on SHAP dimensions and latent vectors. Three SHAP clusters were selected based on clear separation shown by UMAP. The colouring on the UMAP plot is based on the latent space clusters (a-g) and the shape of the marker is based on the SHAP value clustering (α, β, γ). Examining the plot shows that there is no separation between latent clusters c and g, which are grouped together in SHAP cluster γ. Although molecules which belonged to latent clusters a, b, and f were mostly grouped together by SHAP clustering, latent cluster e was grouped together with latent cluster d. Examination of the top dimensions of each SHAP cluster revealed that dimension 24 at least partly encodes for the key sub-structure of clusters a, b, e and f (3,5-pyrazolidinedione, highlighted in dark red), while dimension 26 at least partly encodes for the key sub-structure of cluster d (the oxygen-rich chromenone fused ring system, highlighted in dark green), and dimensions 15, 17, 12 at least partly encode for the key sub-structure of clusters c and g (carboxylic acid bearing aromatic group).

**Figure S16.**
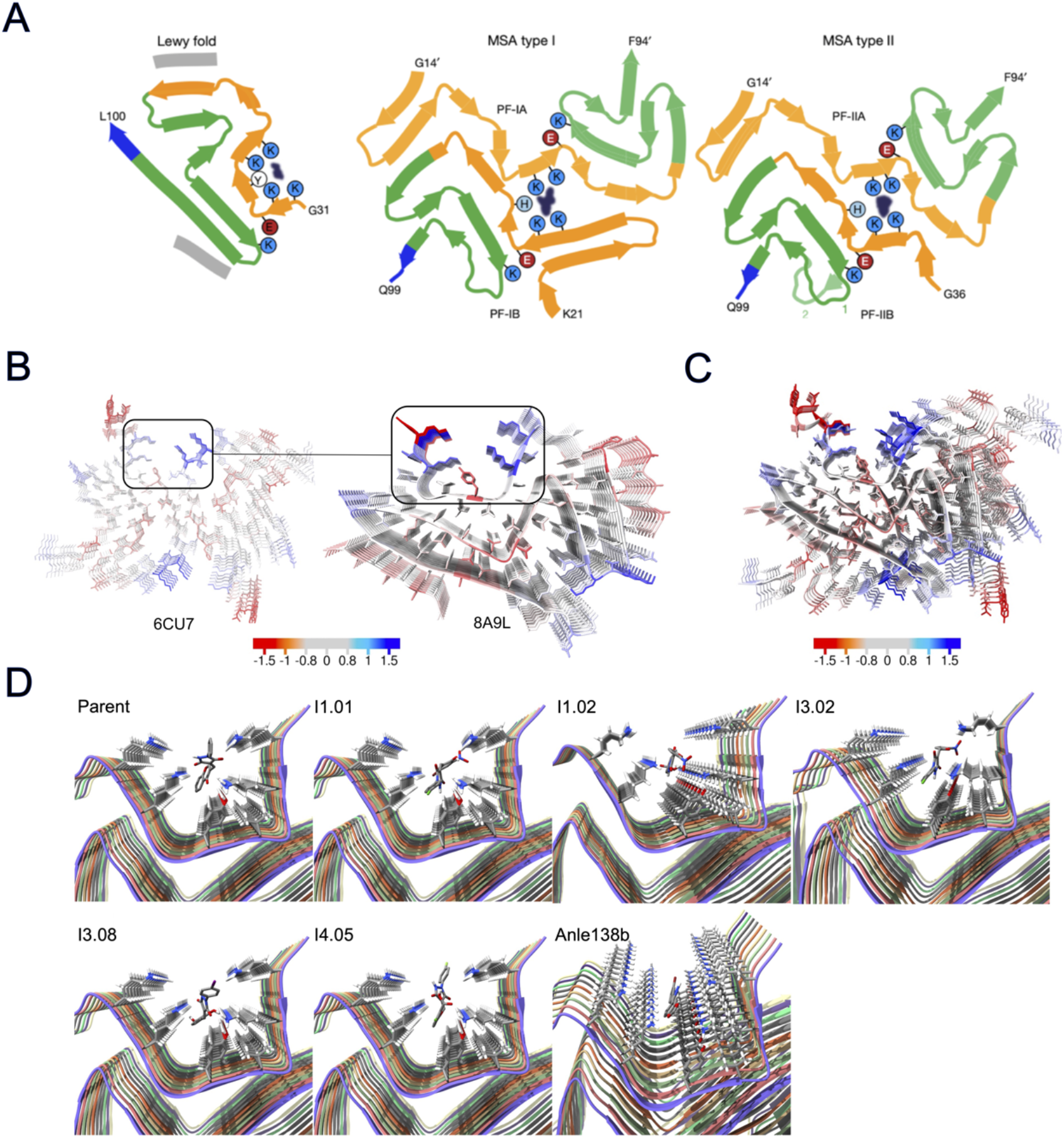
**(A)** Folds of the prevalent fibril polymorph in diseased brain material identified via cryo-EM in Parkinson’s disease and dementia with Lewy bodies (8A9L), MSA type I and MSA type II. A common motif of 4 lysines enclosing an aromatic side chain (tyrosine in the Lewy fold and histidine in the MSA fold and 6CU7 fold) is observed in the polymorphs, with unidentified electron density in the pocket in each case (adapted from Yang, Y. *et al.*^11^). **(B)** Comparison of the cryo-EM structures of the 6CU7 (recombinant, initially targeted) and 8A9L (brain derived) with the homologous binding site indicated. **(C)** Structural overlap of the 6CU7 and 8A9L fibril structures, with the binding site in 6CU7 aligned with the similar binding site in 8A9L at the top of the diagram. The structures are coloured according to the CamSol residue solubility score^6^. **(D)** Schematics of the molecules bound in their lowest energy state within the predicted binding site.

**Figure S17.**
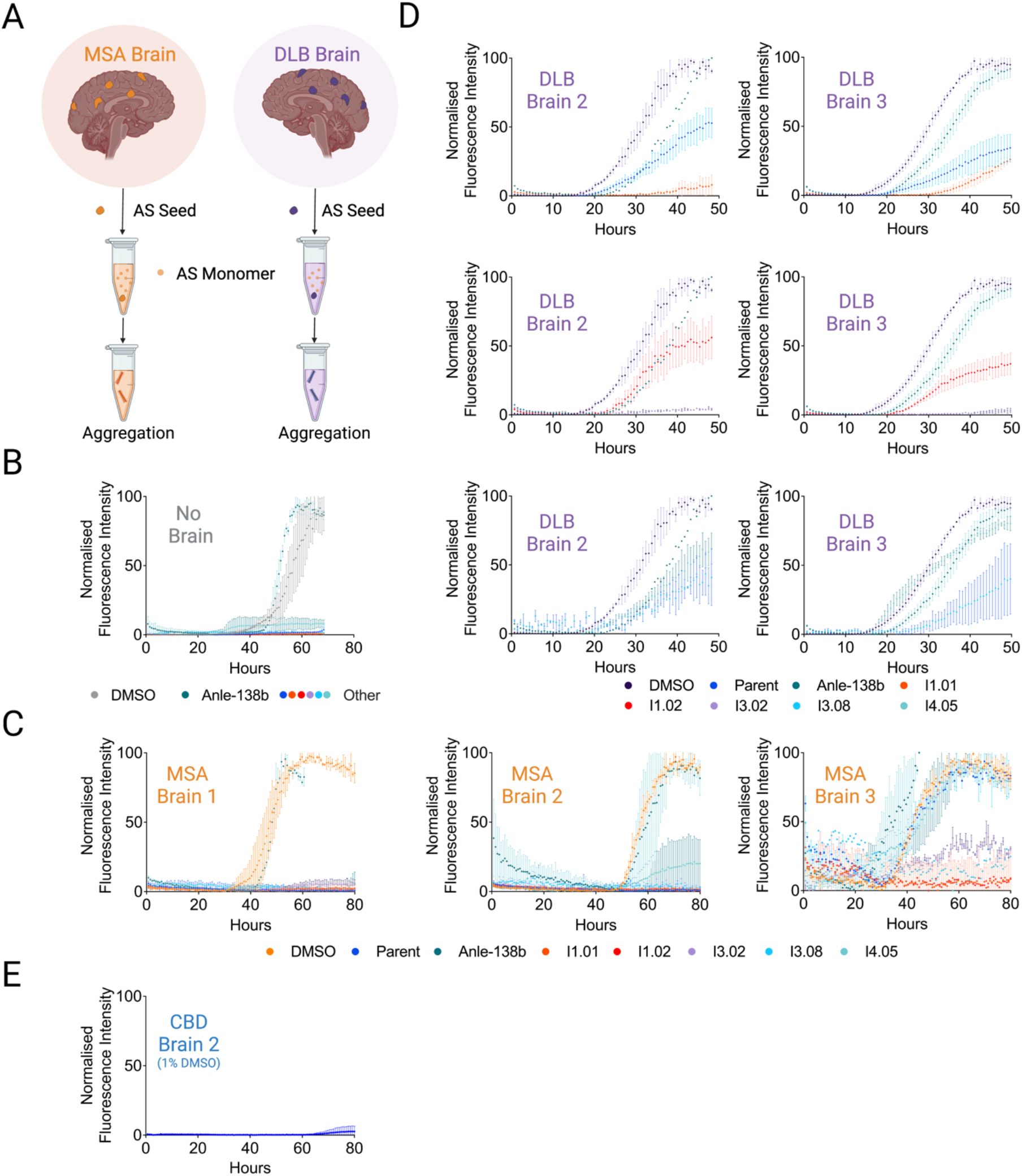
RT-QuIC brain seeding assay. **(A)** Schematic representation of the RT-QuIC assay, aggregates derived from the brain tissue of patients suffering with multiple system atrophy (MSA) or dementia with Lewy bodies (DLB) were used to induce αS aggregation. **(B)** Kinetic traces of a 7 µM solution of αS in the absence of seeds (pH 8, 42°C, shaking at 400 rpm with 1 min intervals, in triplicate, error bars denote SD). Unseeded samples were 1% DMSO (grey), 7 µM Anle-138b (teal), parent (blue), I1.01 (red), I3.02 (lilac), I3.08 (turquoise) and I4.05 (light blue). Anle-138b, in teal, induces aggregation under this condition. **(C)** Kinetic traces of a 7 µM solution of αS in the presence of MSA seeds. The MSA samples were 1% DMSO (light orange), 7 µM Anle-138b (teal), parent (blue), I1.01 (red), I3.02 (lilac), I3.08 (turquoise) and I4.05 (light blue). Anle-138b had no effect in samples 2 and 3 but appears to accelerate aggregation in sample 3. **(D)** Kinetic traces of a 7 µM solution of αS in the presence of DLB seeds. The DLB samples were 1% DMSO (purple), 7 µM Anle-138b (teal), parent (blue), I1.01 (red), I3.02 (lilac), I3.08 (turquoise) and I4.05 (light blue). Data have been separated for clarity. The DMSO and Anle-138b traces are shown on each graph, with 2 molecules from the docking or ML shown for comparison: Parent and I1.01 (top), I1.02 and I3.02 (middle), I3.08 and I4.05 (bottom). Anle-138b exerts a consistent mild inhibition for these two brain samples. **(E)** Kinetic traces of a 7 µM solution of αS in the presence of CBD seeds and 1% DMSO over a longer time course. No significant aggregation was observed over 80 h.

**Figure S18.**
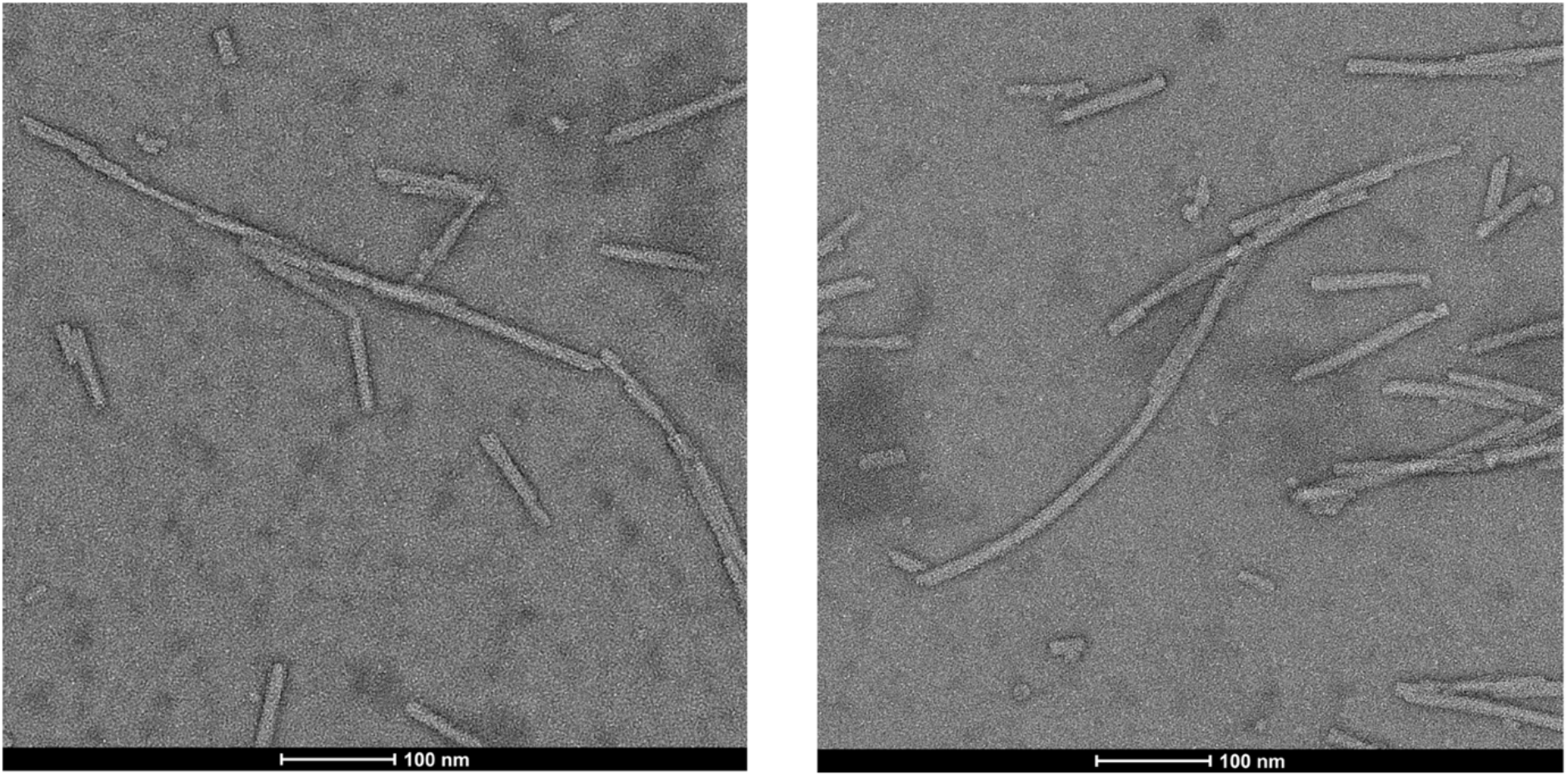
Transmission electron microscopy images of the fibrils at the end of the secondary nucleation assay. Two representative images are shown, the scale bar is 100 nm.

